# Nanoscale visualization of *Drosophila* E-cadherin ectodomain fragments and their interactions using DNA origami nanoblocks

**DOI:** 10.1101/2024.09.02.610731

**Authors:** Hiroki Oda, Shigetaka Nishiguchi, Chihong Song, Kazuyoshi Murata, Takayuki Uchihashi, Yuki Suzuki

## Abstract

The adhesive function of cell surface proteins can be visually assessed through direct observation; however, the underlying structures that mediate adhesion typically remain invisible at the nanoscale level. This hinders knowledge on the diversity of molecular architectures responsible for cell-cell adhesion. *Drosophila* E-cadherin (DE-cadherin), a classical cadherin with a unique domain structure, demonstrates adhesive function; however, it lacks a structural model that explains its adhesion mechanism. In this study, we present a novel application of DNA origami technology to create a cell-free, flat environment in which full DE-cadherin ectodomains are anchored using SNAP-tags and biotin-streptavidin interactions. DNA origami was assembled into a 120 nm long block, bearing 5 or 14 biotin:streptavidin sites that were evenly spaced on one lateral face. DE-cadherin ectodomain fragments were attached via biotinylated SNAP-tags. These decorated DNA origami nanoblocks were subjected to transmission electron and high-speed atomic force microscopy, which revealed a hinge-like site that separated the membrane-distal and -proximal portions of the DE-cadherin ectodomain, suggesting a role in mechanical flexibility. We also observed interactions between DE-cadherin ectodomains via their membrane-distal portions on single DNA origami nanoblocks. We reconstituted an adhesion-like process via pairing DNA origami nanoblocks using DE-cadherin ectodomain interactions. Homophilic associations of functional DE-cadherin ectodomains between the paired DNA origami nanoblocks were visualized at the nanoscale, displaying strand-like molecular configurations, likely representing the extracellular cadherin repeats without regular arrays of structural elements. This study introduces a DNA origami-based platform for reconstituting and visualizing cadherin ectodomain interactions, with potential applications for a broader range of adhesion molecules.

**Highlights:**

- DNA origami technology was applied to perform a structure-function study of cadherin.
- DNA origami nanoblocks decorated with DE-cadherin ectodomains were observed by TEM/HS-AFM.
- A hinge-like site that separated the membrane-distal and -proximal portions of the DE-cadherin ectodomain was revealed.
- An adhesion-like process was mimicked via pairing two nanoblocks using DE-cadherin ectodomain interactions.
- Homophilic associations of DE-cadherin ectodomains between the nanoblocks were visualized at the nanoscale level.

## Introduction

The metazoan genome contains numerous genes that encode cell surface proteins that can mediate cell-cell adhesion (Hynes and Zhao, 2000; Putnam et al., 2007; Srivastava et al., 2010). A major category of cell-cell adhesion molecules is the cadherin superfamily, whose members are involved in various collective cell behaviors, including epithelial organization, pattern formation, morphogenesis, tissue homeostasis, and cancer development (Cavallaro and Christofori, 2004; Janiszewska et al., 2020). The cadherin superfamily is characterized by the presence of repeated extracellular cadherin domains (ECs), each consisting of approximately 110 amino acid residues with specific structural characteristics. However, its members represent considerable diversity in sizes and domain organizations (Nichols et al., 2012; Oda and Takeichi, 2011; Gul et al., 2017). The structural models proposed for adhesion vary among different cadherin types and subtypes (Gul et al., 2017). Ancestral and derived features in cadherin superfamily members have attracted attention for understanding metazoan evolution (Nichols et al., 2012; Oda, 2012).

Classical cadherins, a family found across metazoans but not in non-metazoans, are distinguished by their conserved cytoplasmic domains, which interact with the actin cytoskeleton via catenins and other proteins (Nichols et al., 2012; Oda and Takeichi, 2011; Clarke et al., 2016). These cadherins exhibit Ca^2+^-dependent homophilic adhesion properties through *trans* (between proteins on opposing membranes)*-* and *cis* (between proteins on same membranes)*-*interactions with their ectodomains (Brasch et al., 2012) and act as dynamic clusters at the cell-cell contact sites (Yap et al., 2015). Their crucial role in adherens junction assembly and morphogenesis are conserved across metazoans (Nathaniel Clarke et al., 2019). Despite their functional significance and evolutionary conservation, ectodomains of classical cadherins have diversified and acquired lineage-specific domain organizations through stepwise reductive transitions (Oda et al., 2005; Hulpiau and Van Roy, 2010; Sasaki et al., 2017). Typical classical cadherins in vertebrates possess a derived ectodomain comprising five ECs. Structural studies primarily focus on these 5-EC cadherins, among which the N-terminal-most ECs (EC1s) are major interfaces of *trans-*interactions, mediating adhesion with specificity (Boggon et al., 2002). This 5-EC configuration is conserved in desmosomal cadherins and restricted to vertebrates and urochordates (Gul et al., 2017). In contrast, epithelial classical cadherins in insects, such as *Drosophila* E-cadherin (DE-cadherin), exhibit an independently derived ectodomain with a distinct arrangement of seven ECs, a non-chordate cadherin domain (NC), a cysteine-rich EGF-like domain (CE), and a laminin-G domain (LG) (Oda and Tsukita, 1999; Oda et al., 2005; Sasaki et al., 2017). This cadherin type and other non-chordate classical cadherins lack a structural model that explains its adhesion mechanism. Domain-swapping experiments involving insect E-cadherin ectodomains, however, have identified membrane-distal four ECs, none of which correspond directly to vertebrate EC1 (Oda et al., 2005; Sasaki et al., 2017), as major determinants of homophilic adhesion specificities (Nishiguchi et al., 2016). In contrast to a rod-like configuration of the vertebrate classical cadherin ECs (Pokutta et al., 1994), these four ECs exhibit a folded configuration, as observed via high-speed atomic force microscopy (HS-AFM) (Nishiguchi et al., 2016; Nishiguchi and Oda, 2021). These studies suggest that the core mechanisms of adhesion at adherens junctions fundamentally differ between vertebrates and insects.

The adhesive function of cell surface proteins, such as classical cadherins, can be assessed using aggregation assays using cells that express the protein or microbeads coated with protein fragments (Chappuis-Flament et al., 2001; Lambert et al., 2000; Nagafuchi et al., 1987) Protein-coated glass surfaces (Kovacs et al., 2002), functionalized cantilevers (Baumgartner et al., 2000), protein-anchored liposomes (Lambert et al., 2005), and microfluidic chambers (Eslami Amirabadi et al., 2019) have also been used to study adhesive interactions of cadherin ectodomains. Despite using these techniques and observing tangible functional outcomes, the underlying structures and processes that mediate adhesion remain unknown. Traditional techniques in structural biology, such as X-ray crystallography, cryo-electron microscopy, and HS-AFM, are powerful tools for analyzing cadherin structures and cadherin-cadherin interactions. These methods have been instrumental in modeling classical cadherins and other cadherin superfamily members (Boggon et al., 2002; Maker et al., 2022; Nishiguchi et al., 2022; Cooper et al., 2016). However, visualizing adhesion molecules at the nanoscale level in binding and unbinding states or processes remains challenging because of difficulties associated with reconstructing adhesion assemblies and preparing samples for microscopy. A promising approach involves using liposomes in conjunction with cryo-electron microscopy. Vertebrate classical cadherin ectodomains anchored to liposomes were crystallized to display highly ordered arrays of structural components between opposing liposome membranes, reminiscent of desmosomes (Lambert et al., 2005; Harrison et al., 2011). This method revealed the junction-like outcome; however, the dynamic states of adhesion processes remained inaccessible. Challenges, such as sample thickness and uniformity issues, must also be addressed for wider applicability. The lipid bilayer environment is not essential for *in vitro* reconstruction of classical cadherin ectodomain adhesive interactions (Chappuis-Flament et al., 2001). Involving membrane-free scaffolds may facilitate cooperative interactions of cadherin molecules and help visualize the adhesive and dynamic nature of molecules at the nanoscale level, which could complement the techniques using liposomes.

In this study, we applied DNA origami technology to reconstitute and visualize DE-cadherin ectodomain-adhesive interactions. A key feature of DNA origami is the designability of nanostructures with functionalized sites (Dey et al., 2021). Utilizing this feature, we created a flat environment wherein cadherin ectodomain adhesive interactions occurred and were visualized at the nanoscale level. Transmission electron microscopy (TEM) and HS-AFM was used to visualize DNA origami nanostructures decorated with DE-cadherin ectodomains, which revealed a hinge-like site that separated the membrane-distal and -proximal portions of the DE-cadherin ectodomain, suggesting a role in mechanical flexibility. We mimicked cell-cell interface formation via pairing two DNA origami nanoblocks through DE-cadherin ectodomain interactions. The interfaces between the paired DNA origami nanoblocks showed strand-like molecular structures, likely representing the repeated ECs, although no regular arrays of structural elements were identified. The presented DNA origami-based platform allows the simultaneous realization of adhesion reconstitution and nanoscale visualization, which may have potential applications for structural and functional studies of a broader range of adhesion molecules.

## Results

### Preparing DE-cadherin ectodomain fragments that are biotinylated using a SNAP-tag

We prepared three constructs that each had a whole (denoted DEectW) or partial (denoted DEectD) DE-cadherin ectodomain (Figure 1A). All of them had a V5/6×His-tag at the C-terminal end, and two of them had a SNAP-tag between the DE-cadherin region and the V5/6×His-tag (DEectW-S and DEectD-S; Figure S1). DEectW is identical to the previously described construct DEEXf (Nishiguchi and Oda, 2021). The deleted region in DEectD-S corresponds to EC1-EC4 (Figure 1A; Figure S1), which was previously shown to be a major determinant of homophilic adhesion specificity (Nishiguchi et al., 2016). Using the *Drosophila* S2 cell culture system, we expressed these constructs, which were secreted in the culture media. Proteins were purified from the culture supernatants using cobalt NTA resin. The SNAP-tagged proteins were reacted with biotin-conjugated benzylguanine (BG-bio) (Figure 1B), followed by removal of free BG-bio using dialysis. The resultant biotinylated DEectW-S and DEectD-S products were referred to as DEectW-Sbio and DEectD-Sbio, respectively.

**Figure 1.**
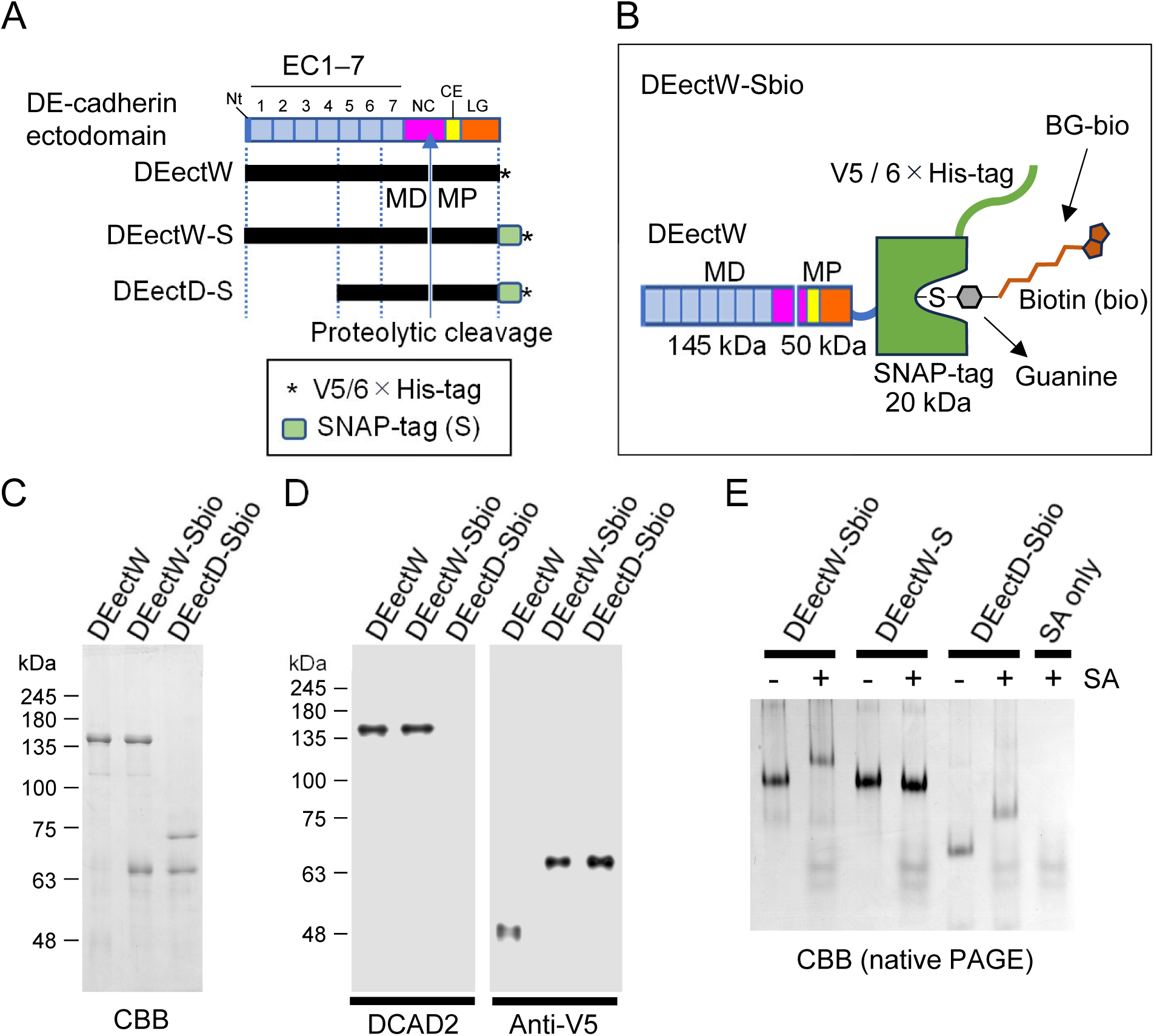
Biotinylated whole or partial *Drosophila* E (DE)-cadherin ectodomain fragment preparation. (A) Schematic representation of the DE-cadherin ectodomain organization, featuring three DE-cadherin ectodomain constructs: DEectW, DEectW-S, and DEectD-S. Nt, N- terminal region; EC, extracellular cadherin domain; NC, non-chordate classical cadherin domain; CE, cysteine-rich EGF-like domain; LG, laminin globular domain. Proteolytic cleavage in the NC domain results in a membrane-distal polypeptide (MD) and a membrane-proximal polypeptide (MP), non-covalently bound to each other (Oda and Tsukita, 1999). (B) Schematic representation of DEectW-S biotinylation via the enzymatic SNAP-tag reaction with BG-biotin, resulting in DEectW-Sbio. (C, D) Coomassie Brilliant Blue (CBB) staining (C) and western blot immunodetection (D) of DEectW, DEectW-Sbio, and DEectD-Sbio separated using sodium dodecyl-sulfate polyacrylamide gel electrophoresis. Approximately 5 pmol of the protein was loaded in each lane in C. The MD fragment was detected with DCAD2, whereas the MP fragment was detected with anti-V5 tag antibody. However, DEectD-Sbio lacked the DCAD2 epitope region (within EC2; (Nishiguchi et al., 2016)). (E) Native PAGE separation and CBB staining of DEectW-Sbio, DEectW-S, and DEectD-Sbio, pre-incubated with streptavidin (SA) at an approximate molar ratio of 1:5 for 10 min, or without SA. Approximately 10 pmol of the cadherin fragment was loaded on each lane.

The matured DE-cadherin ectodomain consists of two polypeptides, resulting from the removal of a prodomain and proteolytic cleavage between Gly^1010^ and Ser^1011^ in the NC, which are referred to as membrane-distal polypeptide (MD) and -proximal polypeptide (MP) (Oda and Tsukita, 1999). Reasonable sizes of the MD and MP for DEectW-Sbio and DEectD-Sbio were observed, as compared with those for DEectW (Figure 1C, D), although no antibody was available for detecting the MD of DEectD- Sbio. Native PAGE mobility shift assay confirmed DEectW-Sbio and DEectD-Sbio could whereas DEectW-S could not bind to streptavidin (SA) (Figure 1E). No traces of non-shifted products were observed for DEectW-Sbio and DEectD-Sbio upon SA addition, indicating a complete reaction of the SNAP-tag with BG-bio.

### Functional characterization of DEectW-Sbio using SA-conjugated microbeads

To test the capability of DEectW-Sbio in mediating homophilic adhesion, we established an aggregation assay system using SA-conjugated microbeads (hereafter referred to as SA beads) (Figure 2A, B). DEectD-Sbio served as a negative control construct in functional assays. SA beads were incubated with 50 nM DEectW-Sbio or DEectD-Sbio for 5 min in HEPES-based Ca^2+^- and Mg^2+^-containing buffer (HCM buffer), followed by adding excess biotin (250 μM) to fill all open biotin-binding sites of the SAs. The beads were then shaken in 4-mm wells at 1,500 rpm for 5 min. After shaking, the microwell was left still for 10 min and the microwell bottom where the beads were sitting was examined by an inverted microscope to determine the degree of bead aggregation, which was defined to be the average size of the top 10% largest bead particles (pixel). Results showed that the degree of bead aggregation clearly differed between DEectW-Sbio and DEectD-Sbio, indicating that DEectW-Sbio could induce bead aggregates, but DEectD-Sbio could not (Figure 2C, D). Two independent modifications, as additional negative controls, were applied to the assay procedure. First, the initial incubation step was performed with neither DEectW-Sbio nor DEectD-Sbio added. Second, SA bead incubation with DEectW-Sbio was preceded by the excess biotin addition step. Neither of these modifications resulted in bead aggregation (Figure 2C, D). Overall, these results suggest that DEectW-Sbio molecules anchored to SA beads via the biotin-SA binding system mediate bead aggregation. Additional time-course analysis showed that the degree of bead aggregation mediated by DEectW-Sbio increased with time during the 5 min of shaking (Figure 2E). The assays performed in the presence of 4 mM CaCl_2_ or 1 mM EGTA indicated that the adhesion capability of DEectW-Sbio was dependent on Ca^2+^ ions (Figure 2F, G). Assays using serially lowered concentrations of DEectW-Sbio (50, 13, and 3 nM) showed that the adhesion activity of DEectW-Sbio was detectable at 50 and 13 nM but not at 3 nM (Figure 2H).

**Figure 2.**
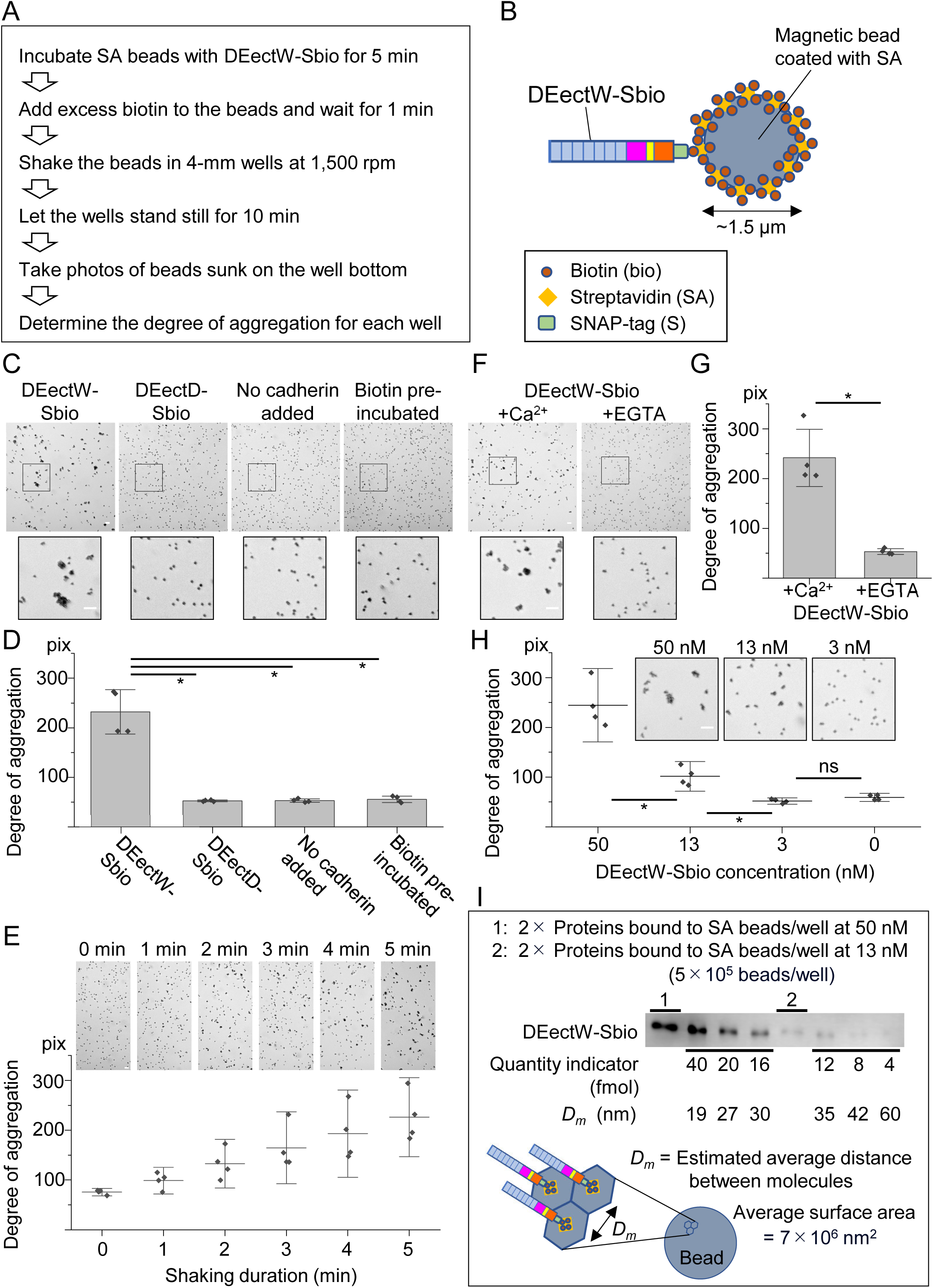
Bead aggregation assays checking for specific adhesive properties of DEectW-Sbio. (A) Outline of the bead aggregation assay procedure. (B) Schematic illustration of an streptavidin (SA)-conjugated microbead capturing the DEectW-Sbio molecule. (C) Images showing aggregate formation of DEectW-Sbio bound microbeads, with corresponding patterns of microbeads obtained for three negative controls, as indicated (DEectD-Sbio used instead of DEectW-Sbio; cadherin fragment omission; SA microbeads pre-incubated with excess biotin used). The boxed region in each panel is magnified below. (D) Quantification of the aggregation degree for assay results shown in C. (E) Images and plots showing increased aggregation of DEectW-Sbio binding microbeads with shaking duration time. (F) Images showing the Ca^2+^-dependency of DEectW-Sbio activity mediating SA microbead aggregation. (G) Quantification of aggregation degree for assay results shown in E. (H) Images and plots showing the concentration dependency of DEectW-Sbio activity. (I) Comparative measurement to assess the DEectW-Sbio protein quantity bound to two aliquots of SA microbeads at concentrations of 50 and 13 nM, with rough estimates of the average distance of DEectW-Sbio molecules on the SA microbead surface. For reference, one aliquot of SA microbeads is defined as the number of microbeads used in each well during aggregation assays, approximately 5 × 10^5^ microbeads. Assuming that the DEectW- Sbio molecules were distributed evenly on the microbead surface, the relationship between the quantity of bound DEectW-Sbio molecules and average distance between the neighboring molecules was calculated. Error bars in D and F indicate standard deviation (s.d.), whereas bars in G and H indicate the mean and 95% confidence intervals. *, p < 0.01 (Welch’s t-test). Scale bars; 10 µm.

To roughly estimate the density of DEectW-Sbio molecules on SA beads, we semi-quantified the number of DEectW-Sbio molecules bound to two aliquots of SA beads (an aliquot of SA beads is 5 × 10^5^ beads, which was used in each microwell in the aggregation assays) in the same conditions as when using DEectW-Sbio at 50 nM and 13 nM in bead aggregation assays (Figure 2I). This quantification showed that each aliquot of SA beads bound ∼20 and ∼6 fmol DEectW-Sbio molecules in the 50 and 13 nM conditions, respectively. Based on the assumption that all DEectW-Sbio molecules were uniformly distributed as monomers on the SA bead surface, the average distance between neighboring DEectW-Sbio molecules was estimated to be ∼19 and 35 nm in the 50 and 13 nM conditions, respectively.

### Constructing and visualizing DNA origami nanoblocks decorated with DEectW- Sbio molecules

To create a flat environment that was conducive to protein fragment anchoring and TEM/HS-AFM observation, we designed a block-shaped, three-dimensional DNA origami nanostructure (∼120 nm × ∼20 nm × ∼10 nm) that had five biotinylated sites at intervals of ∼21.5 nm on one of the long lateral faces (Figure 3A). The DNA origami nanoblock orientation was marked by lacking one of the corners. This DNA origami nanoblock bearing five biotin sites will be hereafter referred to as DO-5×bio. DO-5×bio was decorated with SAs, referred to as DO-5×bio:SA, and these decorated objects were purified using agarose gel electrophoresis (Figure 3B; Figure S2). TEM observation revealed that the nanostructure of DO-5×bio:SA was formed as designed, with one of the largest faces adsorbed onto the support film (Figure 3C).

**Figure 3.**
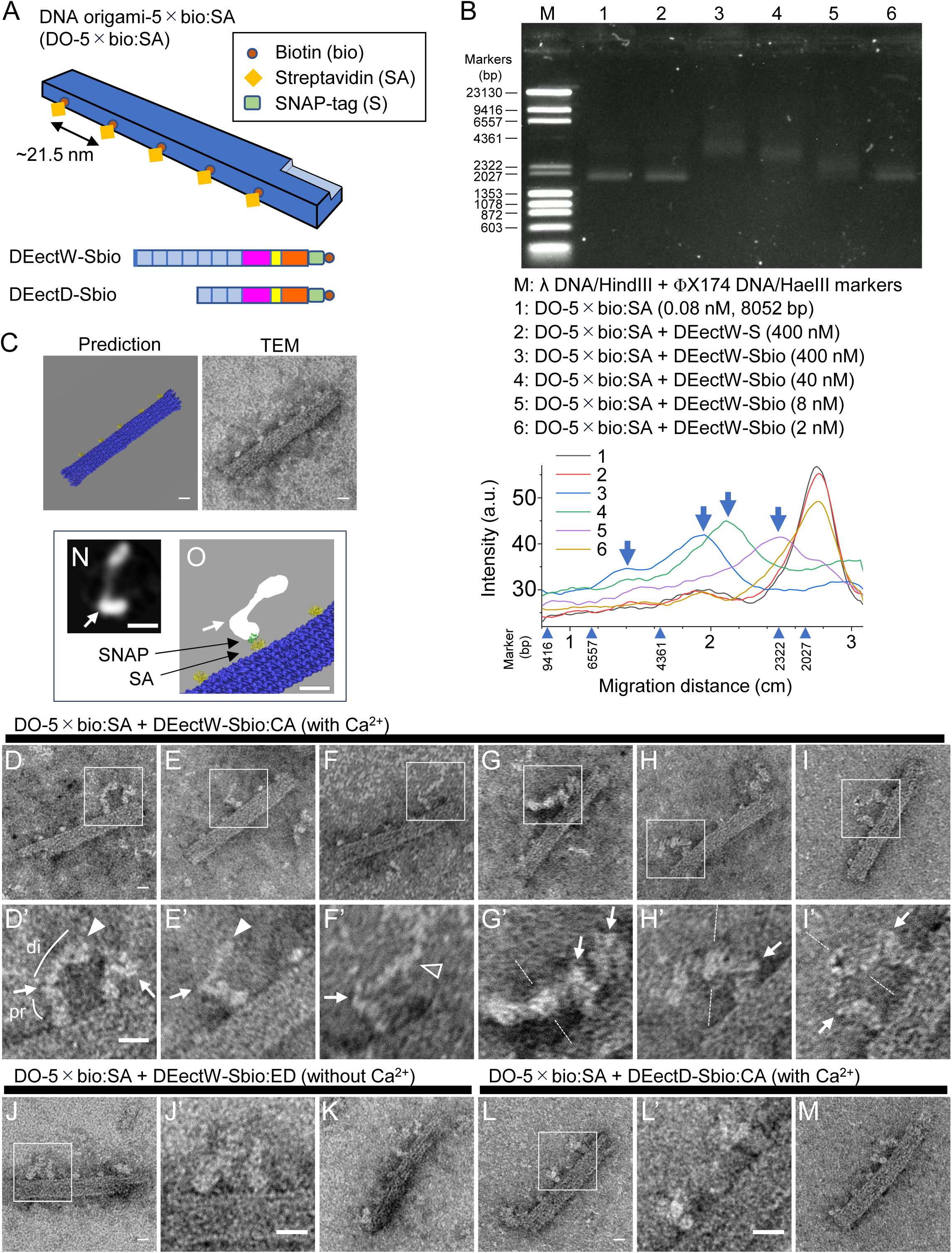
Transmission electron microscopy (TEM) observation of DEectW-Sbio molecules docked onto the DNA origami structure via streptavidin (SA) (A) Schematic representation of the DO-5×bio:SA nanostructure, characterized by a long block shape, an indentation at a corner, and five evenly spaced, biotinylated sites where SA molecules are selectively bound. The designed configuration places these biotinylated sites at intervals of ∼21.5 nm along one long side of the DNA origami. Schematic illustrations of DEectW-Sbio and DEectD-Sbio are also shown. (B) Agarose gel electrophoresis (AGE) analysis checking for DO-5×bio:SA docking with DEectW- SNAP-bio molecules. DO-5×bio:SA was mixed with DectW-S at 400 nM or with DEectW-Sbio at different concentrations (400, 40, 8, and 2 nM) and then incubated for 10 min. DNA migration pattern (left) and DNA stain intensity profiles (right) reveal a concentration-dependent increase in bio:SA site occupation by DEectW-Sbio molecules. (C) Structure prediction of DO-5×bio:SA (left) and TEM observation of negatively stained DO-5×bio:SA (right). (D–M) TEM observation of DO-5×bio:SA nanoblocks onto which DEectW-Sbio (D–K) or DEectD-Sbio (L, M) molecules were docked. The docking was performed in the presence of 4 mM CaCl_2_ (D–I, L, M) or in the absence of Ca^2+^ ions (J, K). Areas boxed in D–I, J, L are magnified in E’–I’, J’, L’, respectively. Arrows pointing to the hinge-like site separating the membrane-distal (di) and -proximal (pr) portions of the *Drosophila* E (DE)-cadherin ectodomain. Closed arrowheads indicate folded configurations of the membrane-distal multiple ECs, whereas open arrowhead indicate an unfolded configuration of the membrane-distal ECs. Broken lines indicate symmetry axes in visualized molecular shapes. (N) A 2D class averaged image of negative-staining TEM images of DEectW molecules from 1,655 particles. The particles were classified into 15 classes and the major class #1 image is presented here (see Figure S3 for details). (O) Proposed structural arrangement of DEectW, SNAP, SA, and DO-5×bio. Scale bars; 10 nm.

We examined the binding of DEectW-Sbio to DO-5×bio:SA. DO-5×bio:SA (final concentration, 0.08 nM) was mixed with DEectW-Sbio at different concentrations (400, 40, 8, and 2 nM) and with 400 nM DEectW-S, followed by 10-min’s incubation and agarose gel electrophoresis. The addition of DEectW-Sbio resulted in differentially shifted peaks of DO-5×bio:SA in a concentration-dependent manner; however, DEectW-S addition did not affect the DO-5×bio:SA migratory behavior (Figure 3B). The peak profiles indicated the presence of at least four states in the binding between DO-5×bio:SA and DEectW-Sbio, which may reflect the multivalency of DO-5×bio:SA. Although DNA origami objects were prepared in Tris-based Folding buffer, which contained 1 mM EDTA (with no Ca^2+^ ions), we ensured that in the Folding buffer supplemented with 4 mM CaCl_2_, DEectW-Sbio exhibited adhesive activity at the same level as that in the HCM buffer (Figure S3).

We used negative staining and TEM to observe DO-5×bio:SA nanoblocks decorated with DEectW-Sbio molecules. DO-5×bio:SA (final concentration, 0.08 nM) was mixed with DEectW-Sbio at an approximate molar ratio of 1:100 in the presence of 4mM CaCl_2_ and then incubated for 2 min, followed by the negative staining procedure. TEM observations identified a total of 70 DO-5×bio:SA blocks in which one or more of the five bio:SA sites were recognizably occupied by DEectW-Sbio molecules (Figure 3D-I), from three independent preparations. These decorated DNA origami nanoblocks in calcium-containing buffer will be referred to as DO-5×bio:SA+DEectW-Sbio:CA (CA denotes Ca^2+^ ions), using the same naming system for all sample types comprising the DNA origami and DE-cadherin construct (Figure S4). Substantial variations were observed in the visualized shape and size of cadherin fragments. However, although each SA on the DNA origami block had theoretically three biotin-binding pockets vacant, only one of the three pockets, which was likely farthest from the DNA origami surface, appeared to have been used for the DEectW-Sbio molecule binding in most cases, possibly because of steric hindrance.

To grasp specific aspects of the structural properties of DEectW-Sbio molecules with Ca^2+^ ions, we examined DEectW-Sbio and DEectD-Sbio molecules in the absence and presence of Ca^2+^ ions, respectively. A total of 49 DNA origami nanoblocks decorated with DEectW-Sbio in EDTA-containing buffer without Ca^2+^ (Figure 3J, J’, K), referred to as DO-5×bio:SA+DEectW-Sbio:ED (ED denotes EDTA), and of a total of 37 DNA origami nanoblocks for DEectD-Sbio with Ca^2+^ (Figure 3L, L’, M), referred to as DO-5×bio:SA+DEectD-Sbio:CA, were identified from two independent preparations for each. Comparisons of visualized cadherin fragments bound to bio:SA sites of the DNA origami nanoblocks confirmed that specifically structured DE-cadherin ectodomain monomers, featured by folded configurations of the membrane-distal multiple ECs, existed in the sample type DO-5×bio:SA+DEectW-Sbio:CA (Figure 3D, D’, E, E’), consistent with our previous observations of DE-cadherin ectodomains using HS-AFM (Nishiguchi et al., 2016; Nishiguchi and Oda, 2021). However, we found exceptions: the membrane-distal ECs were in an unfolded or opened state (Figure 3F, F’). We also found a novel structural feature of the DE-cadherin ectodomain—the presence of a specific hinge-like site that separated the membrane-distal and -proximal portion (Figure 3D’-F’). Flexibility of the hinge angle appeared to allow various bent configurations of the DE-cadherin ectodomain. To understand the flexibility further, we performed two-dimensional (2D) class averaging of the negatively stained TEM images of DEectW molecules, which revealed various shapes of DE-cadherin ectodomain, consistent with the folded, unfolded, and bent configurations of DEectW-Sbio molecules (Figure S5). The negative-staining TEM images (Figure 3D–F) and 2D class averaging of DEectW molecules (Figure 3N) allowed us to propose a structural arrangement for DEectW on SNAP, SA, and DO- 5×bio (Figure 3O), revealing its spatial configuration and potential functional interactions.

Additionally, some molecular images from the sample type DO- 5×bio:SA+DEectW-Sbio:CA displayed potential dimers, exhibiting symmetric shapes (Figure 3G–I, G’–I’). At least one of them indicated an interaction or close proximity of the membrane-distal portions of two DEectW-Sbio molecules rooted at neighboring bio:SA sites (Figure 3I, I’).

### HS-AFM revealed the dynamic nature of the DE-cadherin ectodomain

To examine the structures and dynamics of the DE-cadherin ectodomain on DNA origami scaffold in solution, we visualized DO-5×bio:SA nanostructures decorated with DEectW-Sbio molecules using HS-AFM. DO-5×bio:SA nanoblocks, DEectW-Sbio molecules, and the combined were imaged (Figure 4A–C), showing consistencies with the TEM observations (Figure 3). HS-AFM showed that the membrane-distal and - proximal polarity of the DE-cadherin ectodomain could be recognized (Figure 4B) as the membrane-proximal portion of the DE-cadherin ectodomain was larger in size than that of the folded membrane-distal ECs (Nishiguchi and Oda, 2021). However, when the DEectW-Sbio molecule was anchored to the bio:SA site of the DNA origami nanoblock, the membrane-proximal portion of the DE-cadherin ectodomain was not separately visualized from the bio:SA portion, owing to limited resolution of HS-AFM (Figure 4C). HS-AFM still revealed the dynamics of the membrane-distal portion of DEectW-Sbio molecules. Extending and shrinking dynamics of the DEectW-Sbio molecule were visualized at a time resolution of 0.3 s/frame, showing flexibility mainly derived from the membrane-distal portion of the DEectW-Sbio (Figure 4D and Movie S1). HS-AFM also showed binding and unbinding dynamics of two DEectW-Sbio molecules via their membrane-distal portions on same DNA origami scaffold (Figure 4E and Movie S2). A dimer-like structure formed between a DEectW-Sbio molecule on the DNA origami scaffold and another DEectW-Sbio molecule free from the scaffold were visualized, showing flexibility of the membrane-distal portions of the DEectW- Sbio molecules while retaining binding between protomers (Figure 4F and Movie S3).

**Figure 4.**
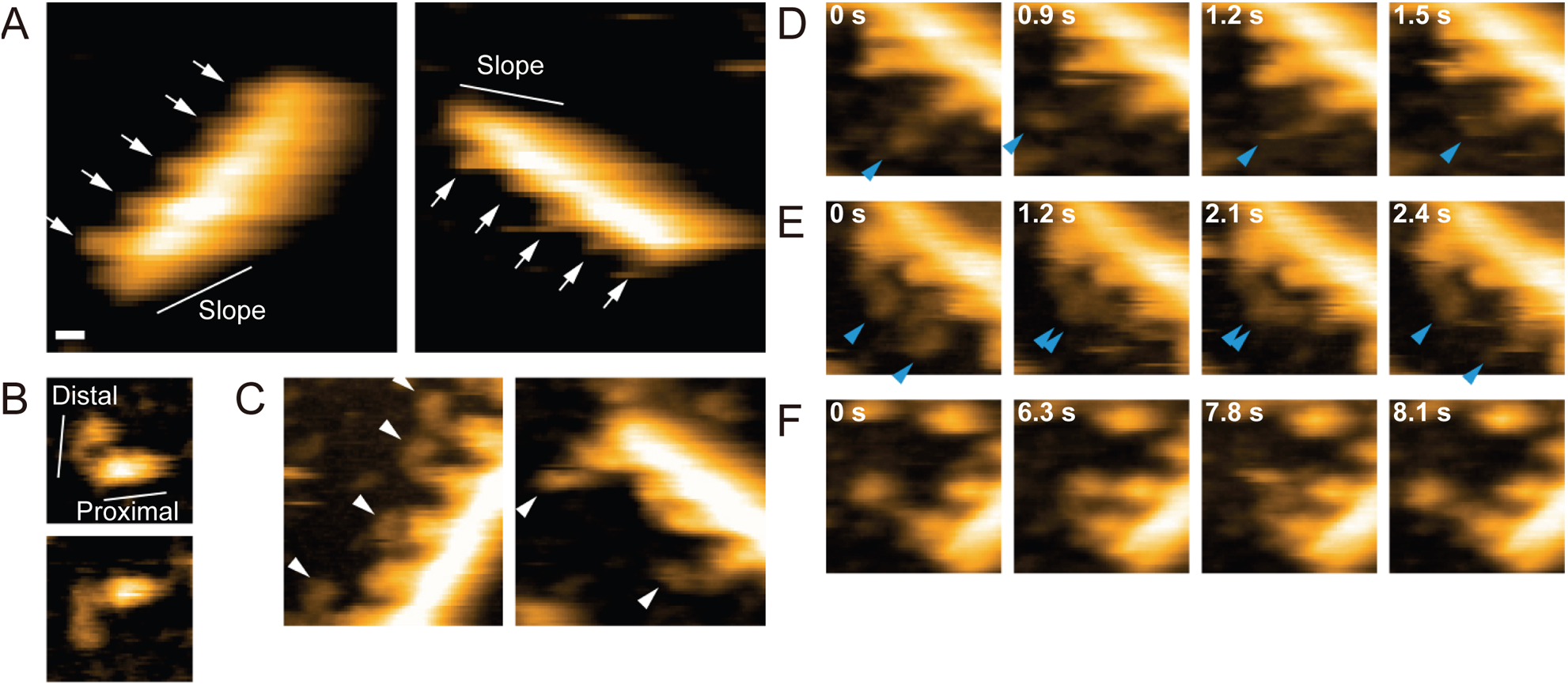
High-speed atomic force microscopy observation of DEectW-Sbio molecules docked onto DNA origami nanoblocks. (A) Two representative images of DO-5×bio:SA structures. Arrows point to the streptavidin (SA)s, which are evenly spaced, on the long side of the rectangle opposite to that with an asymmetric indentation shown with “Slope”. (B) Two representative images of DEectW-Sbio molecules. Membrane-distal and -proximal portions are shown with “Distal” and “Proximal”. (C) Two representative images of DEectW-Sbio docked onto DNA origami nanoblocks. White arrowheads point to the DEectW-Sbio. (D) Sequential images showing extending and shrinking dynamics of membrane-distal potions of the single DEectW-Sbio on the DNA origami nanoblock. Membrane-distal tip region of DEectW-Sbio is indicated by a blue arrowhead. (E) Sequential images showing interactions between membrane-distal portions of two DEectW-Sbio on the same DNA origami nanoblock. Membrane-distal tip regions of two DEectW-Sbio are shown with blue arrowheads. (F) Sequential images showing structural dynamics of the dimer-like structure formed between DEectW-Sbio on DNA origami nanoblock and DNA origami-free DEectW-Sbio. A flexible feature of the membrane-distal portions of DEectW-Sbio with retaining dimer are shown. See Movies S1 to S3. Scale bars; 10 nm.

### Induction and visualization of DE-cadherin ectodomain adhesive interactions between the DNA origami nanoblocks

DE-cadherin ectodomain-mediated adhesion was suggested to depend on the molecule density (Figure 2H, I). To facilitate interactions between DE-cadherin ectodomains on different DNA origami nanoblocks, we modified the DO-5×bio design by replacing nine more staple oligonucleotides with biotinylated ones to produce a DO-14×bio nanoblock, which had two sets of seven biotin sites in line at intervals of ∼7.2 nm on one of the long lateral faces (Figure 5A). DO-14×bio nanoblocks decorated with SAs (DO-14×bio:SA) were examined using negative staining and TEM, confirming the binding of SAs at many, if not all, biotin sites (Figure 5A). Agarose gel electrophoresis showed that DO-14×bio:SA could form a larger complex with DEectW-Sbio molecules than with DO-5×bio:SA (Figure 5B).

**Figure 5.**
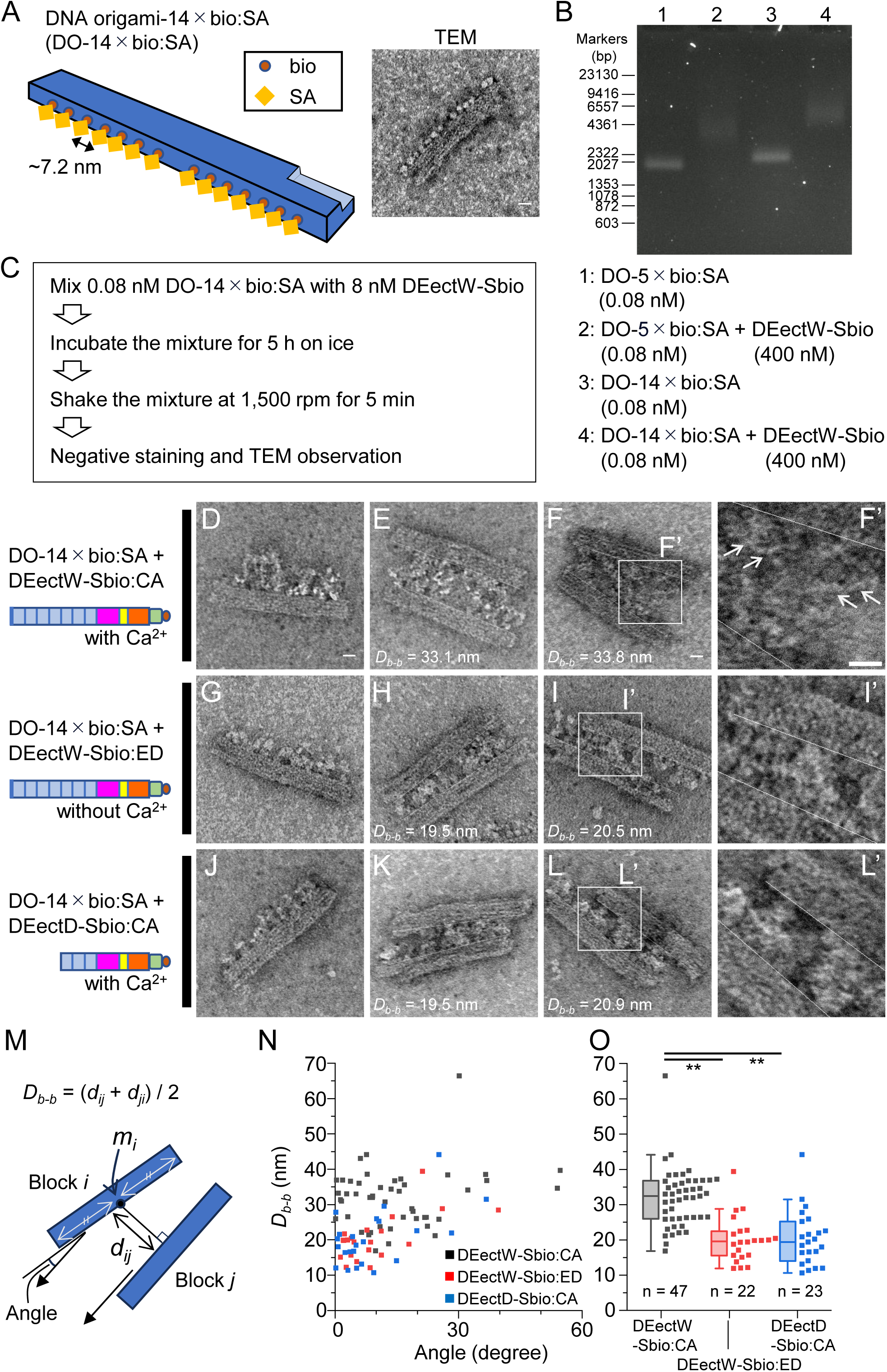
Induction and visualization of *Drosophila* E (DE)-cadherin ectodomain interactions assisted by DNA origami. (A) Schematic representation and transmission electron microscopy image of the DO- 14×bio:SA nanoblock. Two sets of seven evenly spaced bio:SA sites, at intervals of ∼7.2 nm, are placed in a line on a long, lateral face of the block. Bar, 10 nm. (B) Agarose gel electrophoresis analysis comparatively showing DO-5×bio:SA and DO- 14×bio:SA docking with DEectW-Sbio molecules. (C) DNA origami nanoblock pairing procedure. (D–L) Images of single (D, G, J) and paired (E, F, H, I, K, L) DNA origami blocks, formed by combining DO-14×bio:SA with DEectW-Sbio in the presence or absence of Ca^2+^ ions and with DEectD-Sbio in the presence of 4 mM Ca^2+^ ions. Areas boxed in E, H, and K are magnified in E’, H’, and K’, respectively. Bars, 10 nm. (M) Schematic explanation of the defined distance (*D_b-b_*) and angle between paired DNA origami blocks. (N, O) Scatter plots of *D_b-b_* and angle values (N) and box plots of *D_b-b_* values (O) for DNA origami block pairs obtained from the three sample types of the DNA origami pairing are indicated in different colors (gray, DEectW-Sbio:CA; red, DEectW-Sbio:ED; and blue, DEectD:CA).

A mixture of DO-14×bio:SA blocks and DEectW-Sbio molecules was incubated in the presence of Ca^2+^ ions and then shaken, followed by negative staining and TEM (Figure 5C–F). This sample type will be referred to as DO-14×bio:SA+DEectW- Sbio:CA. TEM observations showed that a small proportion of DO-14×bio:SA nanoblocks were found as pairs in which the facing nanoblocks were bridged via DEectW-Sbio molecule interactions (Figure 5E, F); most DO-14×bio:SA blocks remained single, with the bio:SA sites bearing DEectW-Sbio molecules (Figure 5D). A total of 47 DO-14×bio:SA+DEectW-Sbio:CA block pairs were identified and imaged from two independent preparations (https://doi.org/10.6084/m9.figshare.26793652). For comparison, we additionally tested the same combination in the absence of Ca^2+^ ions (DO-14×bio:SA+DEectW-Sbio:ED; Figure 5G–I), and a mixture of DO-14×bio:SA blocks and DEectD-Sbio molecules in the presence of Ca^2+^ ions (DO- 14×bio:SA+DEectD-Sbio:CA; Figure 5J–L). In both sample types, most DO- 14×bio:SA nanoblocks were observed as singles, with the bio:SA sites bearing cadherin fragments (Figure 5G, J). Even in these control sample types, however, paired DO- 14×bio:SA blocks that were bridged through interactions of cadherin fragments were found, with lower frequencies than in the case of DO-14×bio:SA+DEectW-Sbio:CA. A total of 22 DO-14×bio:SA+DEectW-Sbio:ED block pairs and 23 DO- 14×bio:SA+DEectD-Sbio:CA block pairs were identified and imaged from two independent preparations (https://doi.org/10.6084/m9.figshare.26793652).

To compare the dimension of space occupied by the interacting cadherin ectodomain fragments between the three sample types, we measured the distance (*D_b-b_*) and angle between the paired DO-14×bio:SA blocks (Figure 5M), which showed that 75% or more of the DNA origami block pairs for each sample type were at a 20° or smaller angle (Figure 5N). The median *D_b-b_* values for the DO-14×bio:SA+DEectW- Sbio:CA block pairs was 32.5 nm, in contrast to 19.7 and 19.5 nm for the DO- 14×bio:SA+DEectW-Sbio:ED and DO-14×bio:SA+DEectD-Sbio:CA block pairs, respectively (Figure 5N, O). The DO-14×bio:SA+DEectW-Sbio:CA block pairs had larger *D_b-b_* values than the DO-14×bio:SA+DEectW-Sbio:ED and DO- 14×bio:SA+DEectD-Sbio:CA block pairs (Figure 5O), suggesting that homophilic associations of functional DE-cadherin ectodomains were induced and visualized between the facing DNA origami blocks.

Moreover, we examined the morphological aspects of DE-cadherin ectodomain fragments between the facing origami blocks. Strand-like molecular configurations that likely reflected the ECs of DE-cadherin were observed in some of the DO- 14×bio:SA+DEectW-Sbio:CA block pairs (Figure 5F’). Thick strands of protein materials were observed in some of the DO-14×bio:SA+DEectW-Sbio:CA block pairs with the largest *D_b-b_* values (Figure S6), contrasting with the other two sample types (Figure 5I’, L’; Figure S6). However, as the recognized molecular shapes and arrangements varied, interpreting the individual interactions of DE-cadherin ectodomains was difficult. Notably, the observed strand-like structures variously tilted against the surfaces of the paired DNA origami nanoblocks (Figure 5F’).

## Discussion

We applied DNA origami technology to perform a structure-function study of the DE-cadherin ectodomain. The SNAP-tag-biotinylated membrane-proximal side of the DE-cadherin ectodomain showed that it recapitulated Ca^2+^-dependent adhesive properties in a cell and membrane-free environment using SA microbeads, despite the tagging and biotinylation. The strong biotin:SA interactions non-covalently immobilized cadherin fragments to the DNA origami nanoblock at a low concentration, making it compatible with subsequent TEM/AFM observations of molecules without purification. In this application, the tetrameric SA organization may have complicated the anchoring interactions of biotinylated cadherin fragments to the DNA origami object. However, this did not occur based on the peak shift assay of decorated DNA origami (Figure 3B) and TEM/AFM observations of anchored molecules (Figure 3D–I; Figure 4C), which suggested that each bio:SA site on the origami block appeared to serve as a single link to the biotinylated cadherin fragment, presumably because of steric hindrance. The linkage angle was vertical (or near-vertical) to the DNA origami nanoblock surface. These situations facilitate the interpretation of visualized molecule structures and polarities, even when their total shapes vary or change dynamically.

The oriented immobilization of cadherin fragments to the DNA origami nanoblock provides a uniform environment in which adhesion molecules function and are visualized. The designable DNA origami allows immobilized molecules to be constantly placed to be viewed from the lateral side, and their spacing can be easily controlled. The estimated density of DE-cadherin ectodomain fragments in mediating bead aggregation (∼1000–3500 molecules per µm^2^; Figure 2I) was larger, yet roughly comparable to that of DE-cadherin molecules at the adherens junction in epithelial cells of the stage-9 *Drosophila* embryo (∼300 molecules per µm^2^) (Truong Quang et al., 2013). The designed spacing of cadherin fragments on the DNA origami nanoblock was comparable to these functional cadherin densities. Although an apparent demerit of using DNA origami is lack of fluidity in its architecture, this extreme circumstance enables exploring the dynamic nature of cadherin ectodomains, on their own, without considering dynamics of scaffolding materials.

Deploying the DNA origami-based platform has increased our understanding of the structural properties of DE-cadherin ectodomains. A previous study used HS-AFM, in which cadherin fragments that were directly adsorbed on mica substrates were observed; the study suggested that the membrane-distal four ECs of DE-cadherin were folded to form a terminal knot (Nishiguchi et al., 2016). In this study, this structural feature was reproduced not only by HS-AFM but also by TEM. However, as we found some exceptions, the structure of the N-terminal ECs could have multiple states, such as closed and open states (Figure 3F’). This study also discovered that the DE-cadherin ectodomain had a hinge-like linker site between the membrane-proximal and -distal portions. TEM observations captured various hinge angles of this linker site, whereas HS-AFM observations captured extending and shrinking motions of the DE-cadherin ectodomain around the linker site. Overall, these observations suggest that the hinge-like linker site is flexible and facilitates mechanical motions in the DE-cadherin ectodomain, at least in a monomeric state. The precise amino acid residues responsible for the flexible linker site could not be determined because of limitations in resolution of our TEM/HS-AFM measurements. Since some of the morphological features of DE-cadherin ectodomains revealed using negative-staining TEM might be artifacts of the staining and drying procedure, the observed molecular features should be investigated using a more definitive method, such as cryo-electron microscopy.

We observed dimer-like DE-cadherin ectodomains on single DNA origami nanoblocks. Dimers of DE-cadherin ectodomains have not been observed in previous studies using HS-AFM; hence, we considered that the interactions between individual DE-cadherin fragments could be weak (Nishiguchi et al., 2016; Nishiguchi and Oda, 2021). Immobilization of cadherin fragments to the DNA origami nanoblock may affect the thermodynamic stability of interactions, increasing chances for dimer observation. The observed mirror-symmetric shapes of some dimer-like molecules suggested that the DE-cadherin ectodomain dimerization appeared to be mediated by their membrane-distal portions, which were demonstrated to be determinants of homophilic binding specificity (Nishiguchi et al., 2016). HS-AFM directly showed transient and stable interactions between DE-cadherin ectodomains, likely via their membrane-distal portions. Considering the mechanical flexibility of the hinge-like linker site and various motions of the paired molecules, however, we could not determine the mode (*cis-* or *trans*-) of the visualized interaction between the dimer-like molecules.

We reconstituted an adhesion-like process by pairing DNA origami nanoblocks via DE-cadherin ectodomain interactions. Induced adhesion between DNA origami nanoblocks yielded thin, flat architectures that were suitable for TEM sample preparation, providing opportunities to observe homophilic associations of functional DE-cadherin ectodomains at the nanoscale level. The space between the facing DNA origami nanoblocks included two bio:SA layers, which were each ∼5 nm in width (Fan et al., 2019). Considering the distribution of the measured *D_b-b_* values (Figure 5O), the net space available for homophilic associations of DEectW-Sbio molecules between the DNA origami blocks was ∼16–27 nm in width. This space size is comparable to that between the facing cell membranes at the adherens junctions in which adhesion is mediated by DE-cadherin in epithelial cells of *Drosophila* embryos and tissues (Tepass and Hartenstein, 1994). In the presented reconstitution strategy, however, even non-anchored DEectW-Sbio molecules can participate in DE-cadherin ectodomain assemblies forming between the DNA origami nanoblocks. This was particularly probable in DNA origami nanoblock pairs with large *D_b-b_* values (Figure S6A). Based on this situation and strand-like configurations of the likely ECs with no regular arrays of structural elements, DE-cadherin ectodomains may relatively loosely interact with one another to act as protein condensates. Importantly, we showed that DNA origami nanoblock pairing occurred via wild-type DE-cadherin ectodomains without Ca^2+^, as well as via the deleted ones with Ca^2+^, although at lower frequencies. These results could be attributed to pairing DNA origami nanoblocks experiencing much weaker dissociation forces stemming from fluid friction, compared to aggregating microbeads. Therefore, we need a strategy to assess the specificity or strength of adhesive interactions between the DNA origami nanoblocks, as well as to validate the DNA origami-based methods by testing other known molecules.

Unlike cadherin molecules sitting in the lipid bilayer, DEectW-Sbio ectodomains anchored onto the DNA origami nanoblock or magnetic bead may be prevented from behaving flexibly toward minimizing their interaction energies. This restrictive circumstance could have not allowed structurally organized assemblies of DE-cadherin ectodomains to be observed, in contrast to junction-like assemblies of vertebrate cadherin ectodomains formed between liposomes in a previous study (Lambert et al., 2005; Harrison et al., 2011). If the DNA origami nanoblocks are allowed to flexibly slide via setting them on a lipid bilayer (Suzuki et al., 2015), we could observe dynamic processes of adhesion formation and stabilization of DE-cadherin ectodomains using HS-AFM. If the movement of the DNA origami nanoblocks is manipulated by applied external forces, such as using magnetic devices and optical tweezers (Marjoram et al., 2016; Bartsch et al., 2019), we could observe the detaching and/or force-responsive processes of DE-cadherin adhesions. These challenges should be addressed in the future.

DE-cadherin ectodomains anchored onto the DNA origami nanoblock were in a linear spatial arrangement, unlike the microbeads. This circumstance may limit the functional performance of the DE-cadherin ectodomains. DNA origami nanostructures are designable; therefore, the structural design could be improved toward optimizing the performance of the anchored DE-cadherin fragments, which should be based on elucidating the structural mechanisms of DE-cadherin adhesion. DE-cadherin, similar to vertebrate classical cadherins, has been suggested to act as a nanocluster (Cavey et al., 2008; Strale et al., 2015; Truong Quang et al., 2013; Yap et al., 2015). DE-cadherin ectodomains could have multiple binding interfaces or stepwise regulations involved in forming adhesion assemblies. Further development of the DNA origami-based platform could contribute to visualizing such DE-cadherin ectodomain interactions and adhesion processes, as well as to revealing atomic-level structures of functioning and clustered cadherin ectodomains using cryo-electron microscopy.

In conclusion, we applied DNA origami technology for the nanoscale visualization of DE-cadherin ectodomains and their interactions. This technology is designable and can be easily handled; therefore, it has the potential to be applied to various adhesion molecules. For example, the structural, functional, and mechanical aspects of the molecules could be better understood. Homophilic binding and selective sorting properties of adhesion molecules (Nishiguchi et al., 2016) could be beneficial for artificial cell engineering and molecular robotics, which aim to integrate molecular devices into an autonomous system.

## Methods

### DNA construction

To construct the plasmid for expressing cadherin fragments in *Drosophila* S2 cells, we utilized the pAc5.1 V5-His A vector (Thermo Fisher Scientific, MA, USA). The plasmid was designed for DEectW and named as pAcHis-DEectW. It was identical to the previously described pAcHis-DEEXf (Nishiguchi and Oda, 2021). Integrating a DNA fragment that encoded the SNAP-tag (New England Biolabs, MA, USA) into the *Xba*I site of pAcHis-DEectW yielded the newly generated construct, pAcHis-DEectW-S. Two distinct regions of DE-cadherin cDNA, corresponding to amino acid (aa) 1–70 and aa 523–981, were amplified via polymerase chain reaction and seamlessly fused using In-Fusion Cloning technology (Takara Bio, Shiga, Japan) to produce pAcHis-DEectD. The HindIII-BamHI region of pAcHis-DEectD was replaced with that of pAcHis-DEectW-S to produce pAcHis-DEectD-S. Primers used for the plasmid construction are listed in Table S1.

### Preparation of cadherin fragments

*Drosophila* S2 cells, originally obtained from the *Drosophila* community over 2 decades ago (Schneider, 1972), were cultured at 25°C in Schneider’s medium (Thermo Fisher Scientific) and supplemented with 10% heat-inactivated fetal bovine serum (Biowest, Nuaille, France). In comparison to commercially available S2 cells (R69007; Thermo Fisher Scientific), these S2 cells exhibited enhanced cadherin fragment production when transfected with pAcHis-DEEXf-G (Nishiguchi and Oda, 2021). For a standard scale of cadherin fragment expression and purification, S2 cells were seeded at a density of 1.5 × 10^6^ cells/ml in two 90-mm dishes (10 ml medium/dish) approximately 5 h before transfection. In each dish, S2 cells were transfected with 10 µg of plasmid DNA using *Trans*IT-Insect Transfection Reagent (MIR 6100: Mirus Bio, WI, USA). Following 3.5 d of incubation at 25°C, conditioned media were collected from the two dishes and centrifuged at 1,690 g for 15 min at 4°C, resulting in approximately 20 mL of supernatant. This supernatant was concentrated to 1 mL or less using Amicon Ultra-15 Centrifugal Filter, 100 kDa MWCO (UFC910008; Merck KGaA, Darmstadt, Germany). The concentrated supernatant was then diluted by adding Equilibration/Wash buffer (50 mM sodium phosphate pH 7.4, 300 mM NaCl, 10 mM imidazole) to a volume of 10 mL. The resulting solution was applied to 0.5 mL of TALON Metal Affinity Resin (#635501; Takara Bio), washed with Equilibration/Wash buffer, and eluted with Elution buffer (50 mM sodium phosphate pH 7.4, 300 mM NaCl, 150 mM imidazole). The protein concentration of each eluted fraction was measured and peak fractions were combined, followed by concentration to 40 µL or less using vivaspin 500-30K (Cytiva, Uppsala, Sweden). Typical yields of the proteins were 50– 100 µg. For biotinylation, 5 µM purified SNAP-tagged proteins were reacted with 10 µM biotin-conjugated benzylguanine (BG-bio) (S9110S; New England Biolabs) for 30 min at 37°C. The reacted solution was subsequently dialyzed over 3 d using the *Xpress* Micro Dialyzer (MD100, MWCO 12–15 kDa; Scienova, Jena, Germany) to remove free biotin and exchange the buffer (twice with 40 ml of HE buffer [20 mM HEPES pH 7.35, 1 mM EDTA] and four times with 40 ml of HCM buffer [20 mM HEPES pH 7.35, 4 mM CaCl_2_, 10 mM MgCl_2_]). DEectW-Sbio used for docking on DNA origami in the absence of Ca^2+^ was subjected to dialysis using HE buffer over six exchanges. The prepared cadherin fragments were stored at 4°C until use.

### Protein analyses

The size, quantity, and quality of purified proteins were assessed using sodium dodecyl-sulfate polyacrylamide gel electrophoresis (SDS-PAGE, 7.5 or 6% gels) and Coomassie Brilliant Blue staining. For western blotting, the following antibodies were used: rat ant-DE-cadherin antibody (DCAD2 [1:100]; (Oda et al., 1994)), mouse anti-V5 tag antibody (#46-0705 [1:1000]; Thermo Fisher Scientific), and horseradish peroxidase-conjugated anti-rat (NA935V [1:1000]) and anti-mouse (NA931V [1:1000]) IgG antibodies (GE Healthcare, IL, USA). Signals were detected using WesternSure Premium Chemiluminescent Substrate (#926-95010) and the C-DiGit Blot Scanner (LI-COR; NE, USA). Native PAGE, conducted without SDS and a reducing agent, was performed using a 6% gel. Protein quantity was determined using the Qubit Protein Assay Kit and Qubit 2.0 fluorometer (Thermo Fisher Scientific).

### Bead aggregation assays

Magnetic microbeads coated with streptavidin (J-MS-S160S; Medical and Biological Laboratories, Tokyo, Japan) were prepared as a suspension in HCM-BSA buffer (HCM buffer containing 0.01% bovine serum albumin [BSA; A-2143; Sigma Aldrich, MO, USA]) at a density of approximately 5 × 10^5^ beads/µL. For experiments involving Ca^2+^, 1 µL of the SA beads suspension was mixed with 20 µL HCM buffer, followed by a 5- min incubation with either DEectW-Sbio or DEectD-Sbio at appropriate concentrations (50, 13, or 3 nM), after which it was incubated with excess biotin (250 µM; 12987-14; Nacalai Tesque, Kyoto, Japan) for 1 min. Following the SA-masking step, the bead suspension was transferred to a 4-mm well in µ-Slide 15 Well 3D Glass Bottom (#81507; ibidi, Gräfelfing, Germany), which had been pretreated with HCM-BSA buffer and shaken at 1,500 rpm (mixing radius = 1.5 mm) using the Eppendorf ThermoMixer C (Eppendorf, Hamburg, Germany) for 5 min at 25°C. In Ca^2+^-free conditions, HEM-BSA buffer (20 mM HEPES pH 7.35, 1 mM EGTA, 10 mM MgCl_2_, 0.01% BSA) was used throughout the procedure, except for the initial preparation of the DEectW-Sbio stock in HCM buffer for bead aggregation assays. In additional bead aggregation assays, a Tris-based folding buffer (for preparing DNA origami) was used instead of the HEPES-based buffers. After shaking, the multi-well slide was left undisturbed for 10 min. It was then carefully transferred to an inverted microscope (IX71, Evident, Tokyo, Japan), equipped with a cooled CCD camera (CoolSNAP HQ; Roper Scientific, AZ, USA). Three different regions of the well bottom were photographed using a 20× objective lens for each well. The obtained images were then systematically cropped into squares of 800 × 800 pixels, equivalent to a dimension of 256 × 256 µm. These images were analyzed using the ‘Analyze Particles’ function in Image J 1.53t software (National Institutes of Health, MD, USA; Schindelin et al., 2012). During this analysis, areas of individual bead aggregates (pixels) were measured, with a cut-off value of 10 pixels. Measured bead areas were sorted by size to determine the average top 10% bead areas, which were a measure of the aggregation degree (pixel). Each sample/condition type was typically subjected to four independent repetitions of the experiment, utilizing cadherin fragments from at least two separate purifications.

To determine the DEectW-Sbio protein quantity bound to SA microbeads under the same conditions as those used in the aggregation assays, the beads were isolated using centrifugation immediately after incubating the 50 or 13 nM mixture for 5 min. The molecules bound to the beads were then separated via boiling in SDS sample buffer and examined via western blotting to compare with known amounts of DEectW-Sbio using the V5 tag antibody.

### Preparing DNA origami structures

DNA origami nanostructures were designed using the caDNAno2 software (Douglas et al., 2009)(Figure S7; Tables S2 and S3). A total of 239 oligonucleotide species were synthesized (Eurofins Genomics Inc., Tokyo, Japan) and used as staple DNAs for preparing rectangle-shaped origami. To generate the DNA origami-5×bio and −14×bio (DO-5×bio and −14×bio), 5 and 14 specific species of oligonucleotides, respectively, were replaced with corresponding oligonucleotides modified by adding TT-Biotin-TEG when preparing the staple DNA mixtures. The origami unit was assembled in a 20-µL solution containing 10 nM single-stranded, circular scaffold DNA p8064 (tilibit nanosystems, Munich, Germany), 32 nM staple DNAs, 5 mM Tris buffer (pH 8.0), 1 mM EDTA, and 15 mM MgCl_2_. The mixture of scaffold and staple DNAs was annealed by reducing the temperature from 85 to 65°C for over 1 h, followed by a reduction from 65°C to 44°C at a rate of 1°C/h, and finally to 25 °C. After annealing was completed, the origami products were precipitated by adding an equal volume of 15% PEG6000, 5 mM Tris pH 8.0, 1 mM EDTA, and 505 mM NaCl (Stahl et al., 2014). The precipitate was then resolved in 20 µL of Folding buffer (5 mM Tris pH 8.0, 1 mM EDTA, and 15 mM MgCl_2_). For decoration, a 20-µL solution containing biotinylated DNA origami was mixed with 2 µL of 500 mM SA (32243-11; Nacalai Tesque) and incubated for 20 min at room temperature. The decorated DNA origami products were purified using agarose gel electrophoresis and Freeze N’ Squeeze DNA Gel Extraction Spin Columns (#7326165; Bio-Rad Laboratories, Hercules, CA, USA), immediately followed by dialysis with Folding buffer. Electrophoresis of the DNA origami was performed using 0.7 or 1% agarose gel (#161-3106; Bio-Rad Laboratories, CA, USA) and 0.5× Tris/Borate/EDTA buffer containing 5 mM MgCl_2_. To stain the DNA materials in agarose gels, SYBR Gold (S11494; Thermo Fisher Scientific) was used. The DNA origami quantity (8,052 bp) was determined based on the staining intensity relative to the 2,322 bp and 2,027 bp fragments of the lambda DNA/HindIII marker on the agarose gel.

### Docking of cadherin fragments onto DNA origami nanostructures

DEectW-Sbio and DEectD-Sbio, maintained in HCM buffer, were diluted to a concentration of 40 fmol/µL in Folding buffer containing 20 mM CaCl_2_. To prepare DNA origami with cadherin fragments anchored, 4 µL of 1.0 fmol/µl DO-5×bio:SA was combined with 1 µL of the diluted DEectW-Sbio or DEectD-Sbio solution via pipetting in a 1.5 mL Eppendorf tube (#022431081). The mixture was then incubated for 2 min at room temperature, immediately followed by negative staining for TEM observation. To pair DNA origami nanoblocks via cadherin fragments, 4 µL of 0.75–1.0 fmol/µL DO-14×bio:SA was mixed with 1 µL of 40 fmol/µl DEectW-Sbio or DEectD- Sbio via pipetting in a 1.5 mL Eppendorf tube (#022431081) and incubated for 5 h on ice. The mixture was then shaken at 1,500 rpm on the Eppendorf ThermoMixer C (Eppendorf, Hamburg, Germany) for 5 min at 25°C, immediately followed by negative staining for TEM observation. To establish the Ca^2+^-free condition for DEectW-Sbio docking onto DNA origami, DEectW-Sbio that was initially prepared in HE buffer was diluted in Folding buffer, which contained 1 mM EDTA.

### Negative-staining electron microscopy

To observe mixtures of DNA origami and cadherin fragments, the sample was absorbed onto a glow-discharged carbon support film and stained with 2% uranyl acetate. The stained samples were analyzed using an electron microscope (Hitachi H-7600, JOEL JEM-2010). TEM observations for each sample/condition type were conducted using at least two independent preparations of DNA origami objects and cadherin fragments.

For 2D class averaging of DEectW molecules, the sample (2.5□μL) was applied onto a carbon-coated copper grid that was glow-discharged beforehand. After removing the excess sample solution with a filter paper, specimens were stained with a 2% uranyl acetate solution for 30□s. The grids were dried in air after removing the staining solution with a filter paper. EM images were recorded on a DE-20 camera (Direct electron LP) at a nominal magnification of 40,000 using a JEM2200FS electron microscope (JEOL Ltd.) operated at 200□kV accelerating voltage. The energy slit width was adjusted at 20□eV. The image pixel size was 1.4□Å on the camera. A 2– 3□μm under-focused condition was selected to enhance the contrast of EM images. Image analysis was performed with Relion 2.0 (Scheres, 2012) motion correcting the images using the DE_process_frames.py script provided by the manufacturer. A total of 1,655 particles of DEectW were automatically picked from the images after individually correcting the contrast transfer function. The particles were classified at 2D in Relion 2.0.

### HS-AFM

A mica substrate with 1.5 mm diameter and 0.1 mm thickness (Furuuchi Chemical, Tokyo, Japan) was fixed using glue on a glass stage. A 2 μL 0.04% 3- aminopropyltriethoxysilane (APTES) solution was added on a freshly cleaved mica substrate and incubated for 3 min. The APTES-coated mica substrate was washed twice with 80 μL Milli-Q water. A 2 μL DO-5×bio:SA solution (1.0 fmol/µL) was added on the APTES-coated mica substrate for 3 min. A 2 μL DEectW-Sbio solution that was diluted using HCM buffer was then added on the APTES-coated mica substrate for 3 min. The DEectW-Sbio solution concentration was adjusted based on the pilot observation. DO-5×bio:SA and the DEectW-Sbio were individually observed separately using the method described in the previous section. HS-AFM was performed in HCM buffer at room temperature (25°C) using a laboratory-built HS-AFM instrument operated in tapping mode (Ando et al., 2001). BL-AC10DS-A2 cantilevers (Olympus Corporation, Tokyo, Japan) with a length, width, and thickness of 9, 2, and 0.13 μm, respectively, a spring constant of ∼0.1 Nm−1, and resonant frequencies of 400–500 kHz were used in solution. The carbon tip at the end of the cantilever was grown via electron beam deposition and sharpened using argon gas etching to a tip radius of 2–5 nm (Wendel et al., 1995). The free oscillation amplitude of the cantilever was set to 1–2 nm and the amplitude setpoint for feedback control was ∼80% of the free amplitude during scanning. Two or more independent protein purifications, DNA origami nanoblock preparation, and HS-AFM were performed. HS-AFM images were processed using the laboratory-made analysis software FalconViewer based on Igor Pro-8 (WaveMetrics, OR, USA).

### Image processing

Images were processed using Image J, Photoshop 2023, or Illustrator 2023. The structural model of the designed DNA origami block was generated using the Web-based service offered by the Aksimentiev group (https://bionano.physics.illinois.edu/origami-structure). Molecular models were visualized and edited using the open-source version of PyMOL version 2.5.0 (https://www.pymol.org/). Schematic illustrations were created using either PowerPoint (Microsoft, WA, USA) or Illustrator (Adobe, CA, USA). Plots, processed images, and illustrations were assembled using PowerPoint.

### Statistical analysis

OriginPro 2021 (OriginLab) was used for graphing and statistical analyses. One-tailed Welch’s t-test was used to compare the mean of two groups in the bead aggregation assays. One-tailed Mann–Whitney U test was used to compare the mean ranks of two groups for measuring the distance between paired DNA origami blocks.

## Data availability

Raw data on TEM and HS-AFM of DNA origami objects and cadherin fragments obtained in this study are available on figshare (https://doi.org/10.6084/m9.figshare.26793652). Additional data and materials will be available from the authors upon request.

## CrediT author contribution statement

**Hiroki Oda:** Writing-review & editing, Writing-original draft, Funding acquisition, Visualization, Investigation, Methodology, Conceptualization

**Shigetaka Nishiguchi:** Writing-review & editing, Visualization, Investigation, Methodology, Conceptualization

**Chihong Song:** Investigation

**Kazuyoshi Murata:** Investigation, Resources, Funding acquisition **Takayuki Uchihashi:** Methodology, Resources, Funding acquisition **Yuki Suzuki:** Methodology, Investigation, Conceptualization

## Acknowledgments

The TEM work on the DNA origami objects was performed at the “Hanaichi Ultrastructure Research Institute” in Okazaki (Aichi, Japan). We would like to thank Ibuki Kawamata for help with installing caDNAno2; Christian Ganser for technical advice of HS-AFM; and Akiko Noda for technical assistance.

---

**Figure S1. Amino acid sequence and domain composition of DEectW-S**

The entire amino acid sequence of DEectW-S, consisting of 12 defined domains (Pro, NT, EC1 to EC7, NC, CE, and LG) and two tag regions (SNAP [S] and V5/6×His tags). DEectD-S is the same as DEectW-S except that the region indicated by asterisks is missing in DEectD-S. DEectW is the same as DEectW-S except that the SNAP- tag is not present in DEectW. Pro, including signal peptide, is removed for protein maturation. The cleavage in NC occurs between the Gly and Ser residues indicated by “><”.

**Figure S2. Bead aggregation assays using DEectW-Sbio in DNA origami folding buffer without or with Ca^2+^ ions**

Assays were performed in the same way as in Figure 2 except that DNA origami folding buffer (Tris-based, EDTA-containing) was used with (right) or without (left) adding 4 mM CaCl_2_. Error bars indicate standard deviation (s.d.). *, p < 0.01 (Welch’s t-test).

**Figure S3. Agarose gel electrophoresis analysis showing the preparation of DO- 5×bio and its binding to streptavidin**

A 0.7% gel was used.

**Figure S4. Schematic representation of the naming system of sample types comprising the DNA origami and DE-cadherin construct.** Key graphic elements and abbreviations are explained in insets.

**Figure S5. Transmission electron microscopy (TEM) analysis of DEectW molecules**

(A) A TEM image of negative-stained, DEectW molecules. (B) 2D classification using 1,655 particles. Scale bars; 100 nm (A), 20 nm (B)

**Figure S6. Extreme examples of paired DO-14×bio:SA nanoblocks**

(A–C) Transmission electron microscopy images of paired DO-14×bio:SA nanoblocks having the five largest *D_b-b_* values, from each of the three sample types related to Figure 5. (A) DO-14×bio:SA+DEectW-Sbio:CA. Arrows indicate strand-like configurations of *Drosophila* E (DE)-cadherin ectodomain fragments contributing to bridging the apposed DNA origami nanoblocks. (B) DO-14×bio:SA+DEectW-Sbio:ED. (C) DO- 14×bio:SA+DEectD-Sbio:CA.

**Figure S7. DNA origami block design**

Annealing patterns of scaffold (p8064; light blue) and staple DNAs (239 species; arbitrary multiple colors) are shown. Positions of biotinylated sites for DO-5×bio and DO-14×bio are indicated. See Tables S2 and S3 for staple DNA details.

**Movie S1. Extending and shrinking motion of the membrane-distal portion of DEectW-Sbio on DO-5×bio.** The images were captured at a scanning speed of 0.3 s/frame. Image size: 60×60 nm^2^ (60×60 pixels).

**Movie S2. Binding and unbinding of neighboring DEectW-Sbio on DO-5×bio.** The images were captured at a scanning speed of 0.3 s/frame. Image size: 60×60 nm^2^ (60×60 pixels).

**Movie S3. Flexible motion of dimer-like structure formed between DEectW-Sbio on DO-5×bio and DO-5×bio-free DEectW-Sbio.** The images were captured at a scanning speed of 0.3 s/frame. Image size: 60×60 nm^2^ (60×60 pixels).

**Table S1.** Polymerase chain reaction (PCR) primers used for plasmid construction.

**Table S2.** Staple sequences for the DNA origami block.

**Table S3.** Biotinylated staple DNAs.

