## Supplementary material for "Nanoscale visualization of *Drosophila* E-cadherin ectodomain fragments and their interactions using DNA origami nanoblocks": Tables S2-S3

**Table S2.** Staple sequences for the DNA origami block.

| **Name** | **Sequence** | **Replacement** |
| --- | --- | --- |
| 2L-cbd_0[118] | TAACCGTGCAGATTTTGAGTTAGAATAAGAACCACTTTAACG |  |
| 2L-cbd_0[139] | GAGTCTGCCAGTAACAGAGAGAGCGCATCCACCCTAGTAACA |  |
| 2L-cbd_0[160] | CAGTGAGTGGCCAAAGGGTAAAACTGAAGAACCGCAGTTAATGCCCCCT |  |
| 2L-cbd_0[188] | CCAGAATCCTGAGATTCTGACCTAGCATTGTACCCCCCTCAGTCGGAAC |  |
| 2L-cbd_0[223] | GGCCGATGAAAGCGGTCTGGAATTGTATCTTCAAAAACCCTC |  |
| 2L-cbd_0[244] | CAGAGCGTAGGGCGCTATCAGATTGTAAAATATCGGATAAAA |  |
| 2L-cbd_0[265] | ACGTGCTGGTCACGTTTTTGATTAAAATGAGGAAGAGGCTTT |  |
| 2L-cbd_0[286] | TTGCTTTCACACCCTAATGCCTGTTAAAGGATTGCTGCCAGA |  |
| 2L-cbd_0[309] | TTTTTACAGGGCGCGTACTATGG |  |
| 2L-cbd_0[55] | AACTATCAAAACGCCCTTTTTCAATCCATGATATTTTTACCG |  |
| 2L-cbd_0[76] | TTGCCTGACATTTTCAATAGCTTTTGTTGGCAGGTATACATG |  |
| 2L-cbd_0[97] | TTTGATTTGAAATGGAGCAAGAAAATAGCCAGCATAGGAGTG |  |
| 2L-cbd_1[19] | TTTTTATTACCGCCAGGATAGCCAAACAGCTCCTCATTAAAGTTTT |  |
| 2L-cbd_1[35] | CCATTGCGTGCCACGCTGAGAGCCAGTTTT |  |
| 2L-cbd_10[365] | TTTTGGTGGAGCCGCCCGGCCAAGGTGCTG |  |
| 2L-cbd_11[12] | TTTTCCAATCAATAATAGCCTAATTTGCTTTT |  |
| 2L-cbd_12[111] | GCCTTAATACAGAGAAGCCCACGTAGAA |  |
| 2L-cbd_12[132] | CGACTTGACAGGGAATAACCCGGTGGCA |  |
| 2L-cbd_12[153] | GAGGCGTGAGAATTTTGAGCGCGCAAAG |  |
| 2L-cbd_12[195] | AGCCAGCTAATCAGCGGTAATAGAAAATTCATTAAGCAAAATATGGTTT |  |
| 2L-cbd_12[216] | GCCTCAGCAGGAAGGCAAACAAGCCTCA |  |
| 2L-cbd_12[237] | GAGGGGATATTTAAGTCATTGATCGGTT |  |
| 2L-cbd_12[258] | TAACCGTTATTTTGGAGATCTATGACCC |  |
| 2L-cbd_12[279] | TTGGTGTAAATTTTGGAGAGGGGGAGAA |  |
| 2L-cbd_12[300] | ACCGTAAATTTTTTTCTAGCTGCAAGGA |  |
| 2L-cbd_12[307] | CGGATTGTAACGCCAATAAAACACCAGA |  |
| 2L-cbd_12[321] | GTGGGAACCATCAAATCAATAACCCTCA |  |
| 2L-cbd_12[328] | ATTCTCCCACGACGTTGGGAAGGGCTTG |  |
| 2L-cbd_12[342] | TAACAACTGGCCTTGCCGGAGAATGCCT |  |
| 2L-cbd_12[365] | TTTTAACATTAAATGTGAGCGAGCCAGTGCCTCATTACAACTTT |  |
| 2L-cbd_12[48] | CGCTAACTTTATCCAAGAAAAATAACGG |  |
| 2L-cbd_12[69] | TTTTATCAACGATTTATCTTACTGGCAT |  |
| 2L-cbd_12[90] | CTATTTTAAAAATGAAACAATATTACGC |  |
| 2L-cbd_13[119] | CATAAAACGGGAGGATTGAGGTTATTAA |  |
| 2L-cbd_13[12] | TTTTCAGTTACAAAATGAACAAAGTTACTTTT |  |
| 2L-cbd_13[140] | TAGACGGTTTAGCGTATCTTTATTAAAT |  |
| 2L-cbd_13[161] | CACCCTGAATCATAGTCAACAAAGTCAGCAGAGATGAATGGC |  |
| 2L-cbd_13[182] | CGGTTGATTTCCGGTAGATAAGATTTAG |  |
| 2L-cbd_13[196] | AAAAGCCAAGCAAACTCCGGACACCACCAACAGGTCAGGATT |  |
| 2L-cbd_13[224] | AAGCAAACGACGACTGAGTGTCTGTTTG |  |
| 2L-cbd_13[245] | ACGTTAAGCATCTGACAAGAGTTGCCCC |  |
| 2L-cbd_13[266] | TCGCATTAGATGGGACGTGGACAGCAAG |  |
| 2L-cbd_13[287] | TCAGCTCTGGGATAGGCGAAACGCCTGG |  |
| 2L-cbd_13[35] | CATATTAGAGCGTCAATCTAATGATTATCAGATGATGGCATTTT |  |
| 2L-cbd_13[56] | AATAAGACTGAATCGAACCTCGCGGAAT |  |
| 2L-cbd_13[77] | TAACGTCGCACCCACAATCAAACAAAGA |  |
| 2L-cbd_13[98] | CAGCCTTATCAAGAGCAAATCGTAACAT |  |
| 2L-cbd_14[307] | CAACCGTAACCAATTATTATAAATGTTT |  |
| 2L-cbd_14[328] | AATCACCAAATAATAAATCAGCCAATAC |  |
| 2L-cbd_14[365] | TTTTATTCAAAAGGGTGAGAAAGCCTGTAGACGAGAAAAATATT |  |
| 2L-cbd_15[112] | AATACATTTCATTATTCATCGGCATTTTGCCACCA |  |
| 2L-cbd_15[12] | TTTTCAGAAGGAAACCGAGGAAATTAGCAAGGCCGTTTT |  |
| 2L-cbd_15[133] | ACATATAGGTAAATAGCCCCCTTATTAGAGAGCCA |  |
| 2L-cbd_15[154] | ACACCACTCAACCGCATCTTTTCATAATACCCTCA |  |
| 2L-cbd_15[175] | TTGTCACCCAAAGAACCGGAACCAGAGCACCGCCT |  |
| 2L-cbd_15[217] | GAGCATATACAGGCACCTTTAATTGCTCATTCGAG |  |
| 2L-cbd_15[238] | GTACCAAGCATTAATAAGAGGTCATTTTGACTTCA |  |
| 2L-cbd_15[259] | TGTAATATCAATTCGGCTTAGAGCTTAAAGATTAA |  |
| 2L-cbd_15[280] | GCCTTTAGCGCGAGATATAATGCTGTAGGCAAAGC |  |
| 2L-cbd_15[301] | TAAAAATTTAGCTATGTTTTA |  |
| 2L-cbd_15[322] | TATATTTCGCAAATGTACGGT |  |
| 2L-cbd_15[343] | GAGTAATGTTTGACTTCCATA |  |
| 2L-cbd_15[49] | AATACCCTCACCAGAGCAGCACCGTAATTTGGCCT |  |
| 2L-cbd_15[70] | GATTAAGTGGGAATGACAGAATCAAGTTAGGTTGA |  |
| 2L-cbd_15[91] | AGTATGTCCGTCACAGCGTCAGACTGTAGCCGCCG |  |
| 2L-cbd_16[104] | ATTATCATAGCAAAATAATAAGATTATTAACGAAC |  |
| 2L-cbd_16[125] | GAAATTAACATAAAACAAGAACACCAGTACATCGC |  |
| 2L-cbd_16[146] | GAGGGAAAAAGAAACTAATATTAAAAGGGCGCGAA |  |
| 2L-cbd_16[188] | ACCAGCGAATCAATCGTAAAACTGAAAGCGTGGCA |  |
| 2L-cbd_16[209] | AGAATTAGCAATAAAGAGAATAGAAAGGACGGGGA |  |
| 2L-cbd_16[230] | TAAATCAAAGCTAACCTGAGAAAAGGAGGAGCCCC |  |
| 2L-cbd_16[251] | AGTAGTAAAACATTACAAAGGCTGGCAAACTAAAT |  |
| 2L-cbd_16[272] | GGTGGCACTTTTGCGTAGCTACTGCGCGGGGGTCG |  |
| 2L-cbd_16[293] | ATTTGGGTTTCAACGATAAATGCCGCGCATCACCC |  |
| 2L-cbd_16[314] | AACCTGTTTTTAGATGATATT |  |
| 2L-cbd_16[335] | TACATTTTAAATGCACAGTCA |  |
| 2L-cbd_16[372] | TTTTCTGCGAACGAGTAGATTTAGTGTAGGTAAAGTTTT |  |
| 2L-cbd_16[62] | AGCAAAAAAAAGAACCGAAGCTCATGGAGTATTAA |  |
| 2L-cbd_16[83] | AGCCATTACTCCTTGAAATAGGACGCTCAAAACAG |  |
| 2L-cbd_17[19] | TTTTGAAACGTCACCAATGAAACAAATAAA |  |
| 2L-cbd_17[308] | AATATGCCCCTGACAGGAACGCAAACGG |  |
| 2L-cbd_17[329] | GTCTGGAAAATCAATCGCGTCCCGTCGG |  |
| 2L-cbd_17[350] | TAACAGTTCAGAAACCAGCTTTCATCTTTT |  |
| 2L-cbd_18[174] | AGCCGCCCAAAATCCAAAAGGGCGACATGGAATAAGTTTATT |  |
| 2L-cbd_18[209] | AGACCGGCCAAAAAGAAGATCAAGAATAATCGGCAAAATCCCTTTTGAG |  |
| 2L-cbd_18[372] | TTTTGCTTTAAACAGTTGATTCCCAATTTTTT |  |
| 2L-cbd_19[105] | AATAAGTCACCAGAGCGCGTTAAGGTGA |  |
| 2L-cbd_19[119] | GGGTCAGACCAGGCGAATCATCGACAAT |  |
| 2L-cbd_19[126] | TGCCTTGCAGAGCCCGGTCATATTGACG |  |
| 2L-cbd_19[140] | GTGCCCGATTAGCGGCAAGCAAAAGTAA |  |
| 2L-cbd_19[147] | TATAAACCACCCTCCGTTTGCATTGAGG |  |
| 2L-cbd_19[182] | CTATTATGAAAGTAGGCAAAGTTGCCGC |  |
| 2L-cbd_19[189] | TCTGACATAGTAAGAGCAACAAAAAGGACAGGCTGCAAACTTGCTCACA |  |
| 2L-cbd_19[19] | TTTTCCAGAATGGAAAGCGCAGTGCCACCCCTTTCCTCATCCTA |  |
| 2L-cbd_19[210] | CTATCATGCGAACCAGAGAGTAAGGCAA |  |
| 2L-cbd_19[224] | GTTTACCCTAATGCAGGGCGAAGAGACG |  |
| 2L-cbd_19[231] | AGACGACCGTTTTACTTTTGACATCCAA |  |
| 2L-cbd_19[245] | ACCAAAAATTTAGGTTCGCTAAGGGATA |  |
| 2L-cbd_19[252] | TAGCGAGCCCGAAATGCGGATTACTAAT |  |
| 2L-cbd_19[266] | TGCAAAATAGAAAGGAAAGGGGATAGAC |  |
| 2L-cbd_19[273] | GAAGTTTATCAAAATTGCTGACTGAAAA |  |
| 2L-cbd_19[287] | GGGGGTAACGGAACGCGATTACCAGTCC |  |
| 2L-cbd_19[294] | ATAGTAAGTCAGAACTCAACATATTTTC |  |
| 2L-cbd_19[308] | AGACTGGCTACGTTAGGGTTTAAACGTATTTCCAGCCGGGGG |  |
| 2L-cbd_19[315] | ATAGCGTGTCTTTAAACTAAAGGTCAAT |  |
| 2L-cbd_19[329] | TGCGGAACAGGACGTTGTAAACCTCACCGTGCCAGCGGGCCG |  |
| 2L-cbd_19[336] | TCGTCATTGACCATAGTTTCACATTAGA |  |
| 2L-cbd_19[350] | CATTGAAGAACTGGCAAGCTTTCAGATTTT |  |
| 2L-cbd_19[42] | CTCTGAACACAAACCATCGATTAGCACCATTACCACGCAATAGTAAGCA |  |
| 2L-cbd_19[56] | TTCCAGTGTACTCACGGGTATTGAACAA |  |
| 2L-cbd_19[63] | AAGCGTCCAGACGACAGTAGCTAGAGCC |  |
| 2L-cbd_19[77] | GCTTTTGGCCCGGACACTCATTTATCAA |  |
| 2L-cbd_19[84] | ATGATACTGACAGGTGCCTTTCGACTTG |  |
| 2L-cbd_19[98] | TACTGGTGAGAGGGCCGTTTTGCTAATG |  |
| 2L-cbd_2[118] | CATTAAAGAAAGGATTTTGAAATCGTAGGGATAAGGTCACCA |  |
| 2L-cbd_2[139] | CTGATAGTCTAAAAAACCTCCCCCAATAGGGTTTTGCCTGTA |  |
| 2L-cbd_2[160] | TATTAGTCTAACAAAGAACGCTATAGAAAGAGAAGGCCCTCA |  |
| 2L-cbd_2[181] | CAGACAACCGTCAACACCGCTGAAACCATTAAGAGATCTAAA |  |
| 2L-cbd_2[202] | AAGCCGGCGGAATACGTAAAACGTGGCGCGATGAA |  |
| 2L-cbd_2[223] | CGATTTAATAGGGTAGTATCGGTTGGGAAGATACATCTTGAC |  |
| 2L-cbd_2[244] | CGGAACCGTTTGGACCAGTTTGGGCCTCAATACCACATTACC |  |
| 2L-cbd_2[265] | AGGTGCCTTAAAGACGCATCGAGCTGGCATTCATCAAAGCTG |  |
| 2L-cbd_2[286] | AAATCAAGTCAAAGGGTCACGCTGCAAGAACATTAAAGGCTT |  |
| 2L-cbd_2[309] | TTTTACTACGTGAACCTTAATGCGCCGCTTTT |  |
| 2L-cbd_2[55] | CACCGCCCCTTGCTTTACCAACCAAGAAGGAGGTTACCACCC |  |
| 2L-cbd_2[76] | AGGTGAGAAACCCTGCTACAAAGTACCGATAGGTGAGCAAGC |  |
| 2L-cbd_2[97] | CACCAGCTCAGTTGTTAGTTGAAGCAAGTTGATATTGTACCG |  |
| 2L-cbd_20[167] | CTCCTCAGGCTTATCCTGCCGTCTGGGGTATTCTACTAATAG |  |
| 2L-cbd_20[372] | TTTTATGCGATTTTAATCCCCCTCAAATTTTT |  |
| 2L-cbd_21[119] | GTACAAAGTGAGAACAACGCCTTTTCAAAGATTAA |  |
| 2L-cbd_21[140] | GCATTCCACTTTCAGCAGAGGGACAAAGTCAATAG |  |
| 2L-cbd_21[161] | TAGTTAGTGTATGGTAATAAGAGGGTTGTGGGCAATATAAAG |  |
| 2L-cbd_21[182] | GTTTTGTGACGTTACGACATATTATATATTTTTAA |  |
| 2L-cbd_21[19] | TTTTCCCTCAGAACCGCCACCCTAGCCTTT |  |
| 2L-cbd_21[203] | TTCATCAGACCAGGAAAAAGCGTATGAGGTTGCCC |  |
| 2L-cbd_21[224] | AAGAACCAGGACAGTTAGTGAAATCCGCGGTAAAG |  |
| 2L-cbd_21[245] | CAAATCACGAACTGAAACGATATTGCAGCATAAAC |  |
| 2L-cbd_21[266] | CTCATTCGACGGTCCGGCAAAGTGTCCATGTGTTC |  |
| 2L-cbd_21[287] | GCCCTGAACTTAGCTCGTCTCGCAACCAGGCATCA |  |
| 2L-cbd_21[308] | ACGAGTAATCCGCGCCGGCCACCGTCGGTGCAGCC |  |
| 2L-cbd_21[329] | AGATGGTCGCCTGAAACGGAAGCAAGAACCCCTGC |  |
| 2L-cbd_21[350] | AATCATTTACAACGAGAACGTCAGCGTTTT |  |
| 2L-cbd_21[56] | TCATTTTCAAAAAAAATTCTTCGTGTGATTCTGTA |  |
| 2L-cbd_21[77] | CCAATAGTTTTTTCAACGCTCTTAATGGATTAATT |  |
| 2L-cbd_21[98] | TAACACTTAAAGGAATTGAGAATTTCATTTGAAAA |  |
| 2L-cbd_22[372] | TTTTCGCGAAACAAAGGTGAATTACCTTTTTT |  |
| 2L-cbd_23[119] | TTCGGTCGCTGAGGCAGCGGACTACAACGCTCAGT |  |
| 2L-cbd_23[140] | GGAGTTAAAGGCCGCTAAACAACAGACAGATTAGG |  |
| 2L-cbd_23[161] | GGGATCGTCACCCTAATTTTCCGTAACGGCTGAGAGCCTATT |  |
| 2L-cbd_23[189] | AGCATCGGAACGAGGGTAGCAACGGCTAGTGTACAAGAGTAATAACGCC |  |
| 2L-cbd_23[19] | TTTTTTATCAGCTTGCTTTCGAGGTGAATTCAAAAGGCAGAGCCTAGTACC |  |
| 2L-cbd_23[224] | TTTGAGGACTAAAGTTGAAAGGGATATTCATTCAA |  |
| 2L-cbd_23[245] | CATGAGGAAGTTTCAGGGAACACGTAACAGTTGAG |  |
| 2L-cbd_23[266] | CGGGTAAAATACGTAGGCGCAAGTGAATTTACAGG |  |
| 2L-cbd_23[287] | CTACGAAGGCACCACCATGTTCGAGAAACGAACTA |  |
| 2L-cbd_23[308] | ACGAAAGAGGCAAATGTCGAAGTAAATTGAAAAAT |  |
| 2L-cbd_23[329] | ACTAAAACACTCATGTATCATTTAATTTTACCAGT |  |
| 2L-cbd_23[350] | CCCCAGCGATTATACCAAGTTTT |  |
| 2L-cbd_23[56] | CAGCTTGATACCGAAAATCTCCAGGGATTATCACC |  |
| 2L-cbd_23[77] | GCCGACAATGACAAATAATAAGAACCCAAAGTATA |  |
| 2L-cbd_23[98] | CGCCCACGCATAACGAACAACGAGTTTCTGCCGTC |  |
| 2L-cbd_3[105] | AACAGTTAATACCGTACATTGTGTAGCAATACTTC |  |
| 2L-cbd_3[126] | AAGGTTACCCTAAACACACGATCCATCACGCAAAT |  |
| 2L-cbd_3[147] | AGGAGCACTTTAATGACATTCGCCACCGAGTAAAA |  |
| 2L-cbd_3[189] | TACATATAAATCAAGCACTCCCGCCATTATTACGAGGAACAT |  |
| 2L-cbd_3[19] | TTTTCAGCAAATGAAATTTCCAGCGGCTGTTCAGAACCGCCATTTT |  |
| 2L-cbd_3[210] | GCCCGAGGAGCTTGAAGGGAATAAAGGGATTTTAGACAGGAACGGTACG |  |
| 2L-cbd_3[231] | TGTTCCACTAAAGGCGGGCGCGGAGCTAAACAGGA |  |
| 2L-cbd_3[252] | TCCACTAGTAAAGCGTGTAGCTTCCTCGTTAGAAT |  |
| 2L-cbd_3[273] | CTCCAACGTTTTTTTAACCACGACGAGCACGTATA |  |
| 2L-cbd_3[42] | AGCATCATGCAACAAACAGGAGGCCTTGCTGGTAATATCCAGAACAATTTT |  |
| 2L-cbd_3[63] | AAATATCGCGGTCAAATACCTAGTAGAAGAACTCA |  |
| 2L-cbd_3[84] | TATCTGGAGAAGATAATCGTCAGTAATAACATCAC |  |
| 2L-cbd_4[118] | TTTTAAAACAGAAAAAACAACTATTTAATAGAAAGCGATATA |  |
| 2L-cbd_4[139] | CCTTTGCAGATTTTTTCTGTCAATTTAGACAGTTTCTTGCAG |  |
| 2L-cbd_4[167] | ACAAACAATTCGACGATGAATTACCGACGAGCCAGGATTTTGCTTTTGC |  |
| 2L-cbd_4[174] | AGACTTTATTAGAGTATTTTTAGAACCCAGTGTTTTTATAAT |  |
| 2L-cbd_4[181] | AAGTATTCCTTTTACAGCAGTTGTACATGTAAATGCAGCAGCGAAAGAC |  |
| 2L-cbd_4[223] | ATGGTGGACAACATCAGAAACGCGGCCTATGAACGCAGAGGC |  |
| 2L-cbd_4[244] | AGCAGGCTAAAGTGGCTCTCATAAAGTTACCAACTACTTTTT |  |
| 2L-cbd_4[265] | CGGTCCACCTAATGTTTCTCCGCCGTTCAATCATACATTAAA |  |
| 2L-cbd_4[286] | CCCTGAGCATTAATCGGAATTCGTTTTTCGGAACGAATGCCA |  |
| 2L-cbd_4[307] | TGATTGCTGCCCGCCAGCGCCGCAGCCTACCTGCTACCTAAA |  |
| 2L-cbd_4[328] | ACCAGTGACCTGTCGGAAACATCCTCATTAAATTGAGAATAC |  |
| 2L-cbd_4[349] | GCCAGGGAATGAATACGGGAAGACTTGTGAGATTTCTTTGAC |  |
| 2L-cbd_4[372] | TTTTGGCGGTTTGCGTATTGGGC |  |
| 2L-cbd_4[55] | TATCATCGGATTATGAAAAATTTATACAAAGGCTCTCTTAAA |  |
| 2L-cbd_4[76] | AACCACCGAAGGGTCAATAGATAAAGCCACGTTGATAGTTGC |  |
| 2L-cbd_4[97] | TATCATTTCAAAATCAGAACGGGGCTTAATTGCGACAACCAT |  |
| 2L-cbd_5[119] | TAAAGAATCAATTACCTGAGCATAGCTTATATATT | DO-5×bio / DO-14×bio |
| 2L-cbd_5[140] | CAGGTTTCGCAGAGGCGAATTAGAAGAGAACGCGA | DO-14×bio |
| 2L-cbd_5[161] | ATACAGTGAATACCAAGTTACATCAAAAAAATCCA | DO-14×bio |
| 2L-cbd_5[182] | CATCGGGACGGATTCGCCTGAGACTACCACTATAT | DO-5×bio / DO-14×bio |
| 2L-cbd_5[19] | TTTTATTCATCAATATATTTACGGCCTGTTAATTGTATCGGTTTTT |  |
| 2L-cbd_5[224] | ACGAGCCATTCGTAATCATGGGGTAATGCGGGCGC | DO-14×bio |
| 2L-cbd_5[245] | TAAAGCCATCCCCGGGTACCGTGCTCGTGCGCTTT | DO-5×bio / DO-14×bio |
| 2L-cbd_5[266] | AGTGAGCTGAGCCTCCTCACAACACTGGGCATCAG | DO-14×bio |
| 2L-cbd_5[287] | TGCGTTGCGTGCCTGTTCTTCCGTTAACGCTTACG | DO-14×bio |
| 2L-cbd_5[35] | AATCCTGTTTTTTAATGGAAACAGTATTTT |  |
| 2L-cbd_5[56] | ACTTCTGAACAATTTCATTTGACCTTGCTAAATAA | DO-5×bio / DO-14×bio |
| 2L-cbd_5[77] | TAGAACCAAAACAAAATTAATGCTATTATTTGAAA | DO-14×bio |
| 2L-cbd_5[98] | TATTTGCGATGATGAAACAAAAGAATCCCTTCTGA | DO-14×bio |
| 2L-cbd_6[209] | TGTTTCCTGTGTGAAACAATAAGAAATTGTTATCCAAATTTC |  |
| 2L-cbd_6[307] | TTTCTGCGGTTACCTGGTGCC | DO-5×bio / DO-14×bio |
| 2L-cbd_6[328] | TTTTCACCCGGTGCTGCCAAC | DO-14×bio |
| 2L-cbd_6[349] | CGGCCAGGATCCAGGGTCAGC | DO-14×bio |
| 2L-cbd_6[372] | TTTTTGTGCACTCTGTCGCGCGGGGAGATTTT |  |
| 2L-cbd_7[105] | CATAGCGAAAAGAAACGTAAAAGTTTGA |  |
| 2L-cbd_7[126] | GACGCTGATTCATTATTGCGTCCGAACG |  |
| 2L-cbd_7[147] | TGAATTTAAAATCGAACGTCAAACTCGT |  |
| 2L-cbd_7[189] | CCTCCGGCTTCACTCCGGGTAGGTTGGGAAAAAATTGCTCATCGCCATT |  |
| 2L-cbd_7[19] | TTTTCATAAATCAATAATAAACACCGGAATCATAATTTT |  |
| 2L-cbd_7[210] | TGCGGCTTCATAGCATTCCACTTCCGAA |  |
| 2L-cbd_7[231] | GTTTCTTAGCTCGAGGAAGCAGAAAATC |  |
| 2L-cbd_7[252] | ATCCCTTGTTGAGGTGGGGTGCGCTGGT |  |
| 2L-cbd_7[273] | AGCAAATGCGTCCGTAACTCAAGAGTTG |  |
| 2L-cbd_7[294] | GATGCCGCAGCACGCGCTCACCCTTCACAACCGTCTATCATTTT |  |
| 2L-cbd_7[315] | AGCGGTGGGTCATATCGGGAAAGACGGGCAACAGC |  |
| 2L-cbd_7[336] | ATCAGACAATGCGGCTGCATTTGGTTTTTCTTTTC |  |
| 2L-cbd_7[63] | AATCGTCTACATTTAATAATGAGAAGGA |  |
| 2L-cbd_7[84] | TTCCCTTCATCAAGTACCATATTGCGGA |  |
| 2L-cbd_8[111] | TTAGTTAATCGCCAATGTTCATATTTTC |  |
| 2L-cbd_8[132] | GAAAACTAACATGTCAGACGATACCGCG |  |
| 2L-cbd_8[153] | ATCGCAACATTTTCAAAAGGTAATCAGA |  |
| 2L-cbd_8[174] | GTAAATGCTGATGCTCATAGGTCTGAGATTGCTTTAACAGTA |  |
| 2L-cbd_8[216] | GGTTGCGCGCACAGAGCGGATCGCAACT |  |
| 2L-cbd_8[237] | CGCACTCTGAAGGGCGGAAAATCGGTGC |  |
| 2L-cbd_8[258] | CGGGGTCGCTGATTGTGGTGATTACGCC |  |
| 2L-cbd_8[279] | GCTGGAGCGCGGTCTGTGAGAGGATGTG |  |
| 2L-cbd_8[300] | ATCCCACGTCGCTGATGTTTAAGTTGGG |  |
| 2L-cbd_8[321] | GGCAGCAGAGCACAATCGGCGTCCCAGT |  |
| 2L-cbd_8[342] | AGCAACCCGTGCCGCGGATAAACGACGG |  |
| 2L-cbd_8[365] | TTTTTGGTGCTGGTCTCGCAGTGTCACTGCGCGCCTTTT |  |
| 2L-cbd_8[48] | GGCGTTAATATGCGAATATCCTATCATT |  |
| 2L-cbd_8[69] | TACCGACACCAGTATAAGTCCTAAACCA |  |
| 2L-cbd_8[90] | CCTAAATAACAGTACGCCTGTCGAGAAC |  |
| 2L-cbd_9[12] | TTTTTTACTAGAAAAAAGCATGTAGAAATTTT |  |
| 2L-cbd_9[196] | CCCGTAACGCATAGGCTTCCACGTCTTGGCTGACC |  |
| 2L-cbd_9[35] | TAGTATCAATAAGATATGTGAGTGAATAAATTACCATTGTTTATATTCC |  |

**Table S3.** Biotinylated staple DNAs

| **Name** | **Sequence** | **Use** |
| --- | --- | --- |
| 2L-cbd_5[56]_2T-bio | ACTTCTGAACAATTTCATTTGACCTTGCTAAATAATT-[Biotin-TEG] | DO-5×bio / DO-14×bio |
| 2L-cbd_5[119]_2T-bio | TAAAGAATCAATTACCTGAGCATAGCTTATATATTTT-[Biotin-TEG] | DO-5×bio / DO-14×bio |
| 2L-cbd_5[182]_2T-bio | CATCGGGACGGATTCGCCTGAGACTACCACTATATTT-[Biotin-TEG] | DO-5×bio / DO-14×bio |
| 2L-cbd_5[245]_2T-bio | TAAAGCCATCCCCGGGTACCGTGCTCGTGCGCTTTTT-[Biotin-TEG] | DO-5×bio / DO-14×bio |
| 2L-cbd_6[307]_2T-bio | TTTCTGCGGTTACCTGGTGCCTT-[Biotin-ON] | DO-5×bio / DO-14×bio |
| 2L-cbd_5[77]_2T-bio | TAGAACCAAAACAAAATTAATGCTATTATTTGAAATT-[Biotin-TEG] | DO-14×bio |
| 2L-cbd_5[98]_2T-bio | TATTTGCGATGATGAAACAAAAGAATCCCTTCTGATT-[Biotin-TEG] | DO-14×bio |
| 2L-cbd_5[140]_2T-bio | CAGGTTTCGCAGAGGCGAATTAGAAGAGAACGCGATT-[Biotin-TEG] | DO-14×bio |
| 2L-cbd_5[161]_2T-bio | ATACAGTGAATACCAAGTTACATCAAAAAAATCCATT-[Biotin-TEG] | DO-14×bio |
| 2L-cbd_5[224]_2T-bio | ACGAGCCATTCGTAATCATGGGGTAATGCGGGCGCTT-[Biotin-TEG] | DO-14×bio |
| 2L-cbd_5[266]_2T-bio | AGTGAGCTGAGCCTCCTCACAACACTGGGCATCAGTT-[Biotin-TEG] | DO-14×bio |
| 2L-cbd_5[287]_2T-bio | TGCGTTGCGTGCCTGTTCTTCCGTTAACGCTTACGTT-[Biotin-TEG] | DO-14×bio |
| 2L-cbd_6[328]_2T-bio | TTTTCACCCGGTGCTGCCAACTT-[Biotin-TEG] | DO-14×bio |
| 2L-cbd_6[349]_2T-bio | CGGCCAGGATCCAGGGTCAGCTT-[Biotin-TEG] | DO-14×bio |
