## Supplementary Information for "Nanoscale visualization of *Drosophila* E-cadherin ectodomain fragments and their interactions using DNA origami nanoblocks"

Contents:

**Figure S1. Amino acid sequence and domain composition of DEectW-S.**

**Figure S2. Agarose gel electrophoresis analysis showing the preparation of DO-5×bio and its binding to streptavidin.**

**Figure S3. Bead aggregation assays using DEectW-Sbio in DNA origami folding buffer without or with Ca^2+^ ions.**

**Figure S4. Schematic representation of the naming system of sample types comprising the DNA origami and DE-cadherin construct.**

**Figure S5. Transmission electron microscopy (TEM) analysis of DEectW molecules.**

**Figure S6. Extreme examples of paired DO-14×bio:SA nanoblocks.**

**Figure S7. DNA origami block design.**

**Legends of Movies S1-S3.**

Note that Table S1, Tables S2 and S3 are provided in separate files.

Prodomain (Pro: this domain is removed for protein maturation)

```
-69..-1(69)
MSTSVQRMSRSYHCINMSATPQAGHLNPAQQQTHQQHKRKCRDLGRRLIPARLLLGVIVAISLLSPALA
N-terminal domain (NT)
1..14(14)
LHSPPDKNFSGDNR
 *************
EC1
15..122(108)
KPAFKNCAGYAPKVKEEQPENTYVLTVEAVDPDPDQVIRYSIVQSPFERPKFFINPSTGVIFTTHTFDRDEPIHEKFVFVTVQATDNGLPPLDDVCTFNVTIEDINDN
************************************************************************************************************
EC2
123..228(106)
APAFNKARYDESMSENAQPDAVVMTISASDFDDGNNSLVEYEILRERDFQYFKIDKESGIIYLKRPIDKRPGQSYAIIVRAYNVVPDPPQDAQIEVRIRVVESSIK
**********************************************************************************************************
EC3
229..339(111)
PPSFVNPIDTPIYLKENLKNFTHPIATLRAVSNMPDKPEVIFELNTGRTEQTNSKNTFVFNQIGNEVTISLGKTLDYEAITDYTLTMIVRNTHELGTEHQIKIQVEDVNDN
***************************************************************************************************************
EC4
340..449(110)
IPYYTEVKSGTILENEPPGTPVMQVRAFDMDGTSANNIVSFELADNREYFTIDPNTGNITALTTFDREERDFYNVKVIASDNSPSSLFDNGEPNRGHQVFRISIGDKNDH
**************************************************************************************************************
EC5
450..550(101)
KPHFQQDKYLAERLLEDANTNTEVIEVKAEDEDNASQILYSIESGNVGDAFKIGLKTGKITVNQKLDYETITEYELKVRAFDGIYDDYTTVVIKIEDVNDN
****
EC6
551..660(110)
PPVFKQDYSVTILEETTYDDCILTVEAYDPDIKDRNADQHIVYSIHQNDGNRWTIDNSGCLRLVKTLDRDPPNGHKNWQVLIKANDEDGVGTTVSTVKEVTVTLKDINDN
EC7
661..765(105)
APFLINEMPVYWQENRNPGHVVQLQANDYDDTPGAGNFTFGIDSEATPDIKTKFSMDGDYLHANVQFDREAQKEYFIPIRISDSGVPRQSAVSILHLVIGDVNDN
Nonchordate classical cadherin domain (NC)
766..976(211)
AMSEGSSRIFIYNYKGEAPETDIGRVFVDDLDDWDLEDKYFEWKDLPHDQFRLNPSTGMITMLVHTAEGEYDLSFVVTEDSMFVPRHSVDAYVTVVVRELPEEAVDKSGSIRFINVTKEEFISVPRDFQSPDALSLKDRFQLSLAKLFNTSVSNVDVFTVLQNENHTLDVRFSAHG><SPYYAPEKLNGIVAQNQQRLENELDLQMLMVNIDE
Cysteine-rich EGF-like domain (CE)
977..1055(79)
CLIEKFKCEESCTNELHKSSVPYMIYSNTTSFVGVNAFVQAQCVCEAPLMRRCLNGGSPRYGENDVCDCIDGFTGPHCE
Laminin Globular domain (LG)
1056..1247(192)
LVSVAFYGSGYAFYEPIAACNNTKISLEITPQIDQGLIMYLGPLNFNPLLAISDFLALELDNGYPVLTVDYGSGAIRIRHQHIKMVADRTYQLDIILQRTSIEMTVDNCRLSTCQTLGAPIGPNEFLNVNAPLQLGGTPVDLEQLGRQLNWTHVPNQKGFFGCIRNLTINEQTYNLGMPSVFRNIDSGCQQS
SNAP-tag#
1248..1431(184)
SRMDKDCEMKRTTLDSPLGKLELSGCEQGLHRIIFLGKGTSAADAVEVPAPAAVLGGPEPLMQATAWLNAYFHQPEAIEEFPVPALHHPVFQQESFTRQVLWKLLKVVKFGEVISYSHLAALAGNPAATAAVKTALSGNPVPILIPCHRVVQGDLDVGGYEGGLAVKEWLLAHEGHRLGKPGLG
V5/6xHis-tag
1432..1460(38)
SRGPFEGKPIPNPLLGLDSTRTGHHHHHH
```

**Figure S1. Amino acid sequence and domain composition of DEectW-S.** The entire amino acid sequence of DEectW-S, consisting of 12 defined domains (Pro, NT, EC1 to EC7, NC, CE, and LG) and two tag regions (SNAP [S]- and V5/6×His-tags). DEectD-S is the same as DEectW-S except that the region indicated by asterisks is missing in DEectD-S. DEectW is the same as DEectW-S except that the SNAP- tag is not present in DEectW. Pro, including signal peptide, is removed for protein maturation. The cleavage in NC occurs between the Gly and Ser residues indicated by “><”.

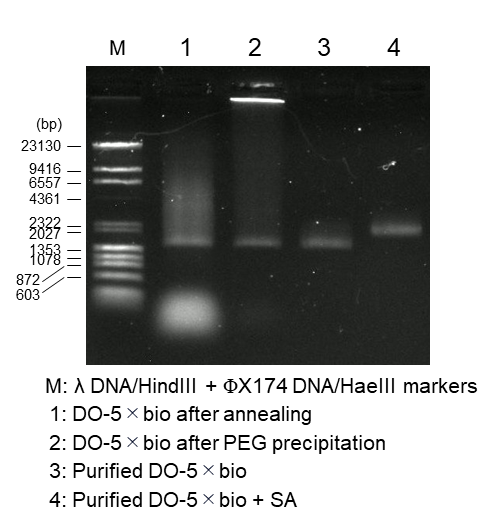

**Figure S2. Agarose gel electrophoresis analysis showing the preparation of DO-5×bio and its binding to streptavidin (SA).** A 0.7% gel was used.

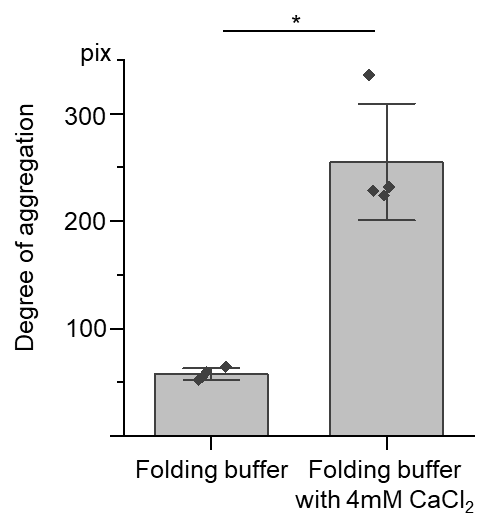

**Figure S3. Bead aggregation assays using DEectW-Sbio in DNA origami folding buffer without or with Ca^2+^ ions.** Assays were performed in the same way as in Fig. 1 except that DNA origami folding buffer (Tris-based, EDTA-containing) was used with (right) or without (left) adding 4 mM CaCl_2_. Error bars indicate standard deviation (s.d.). *, p < 0.01 (Welch’s t-test).

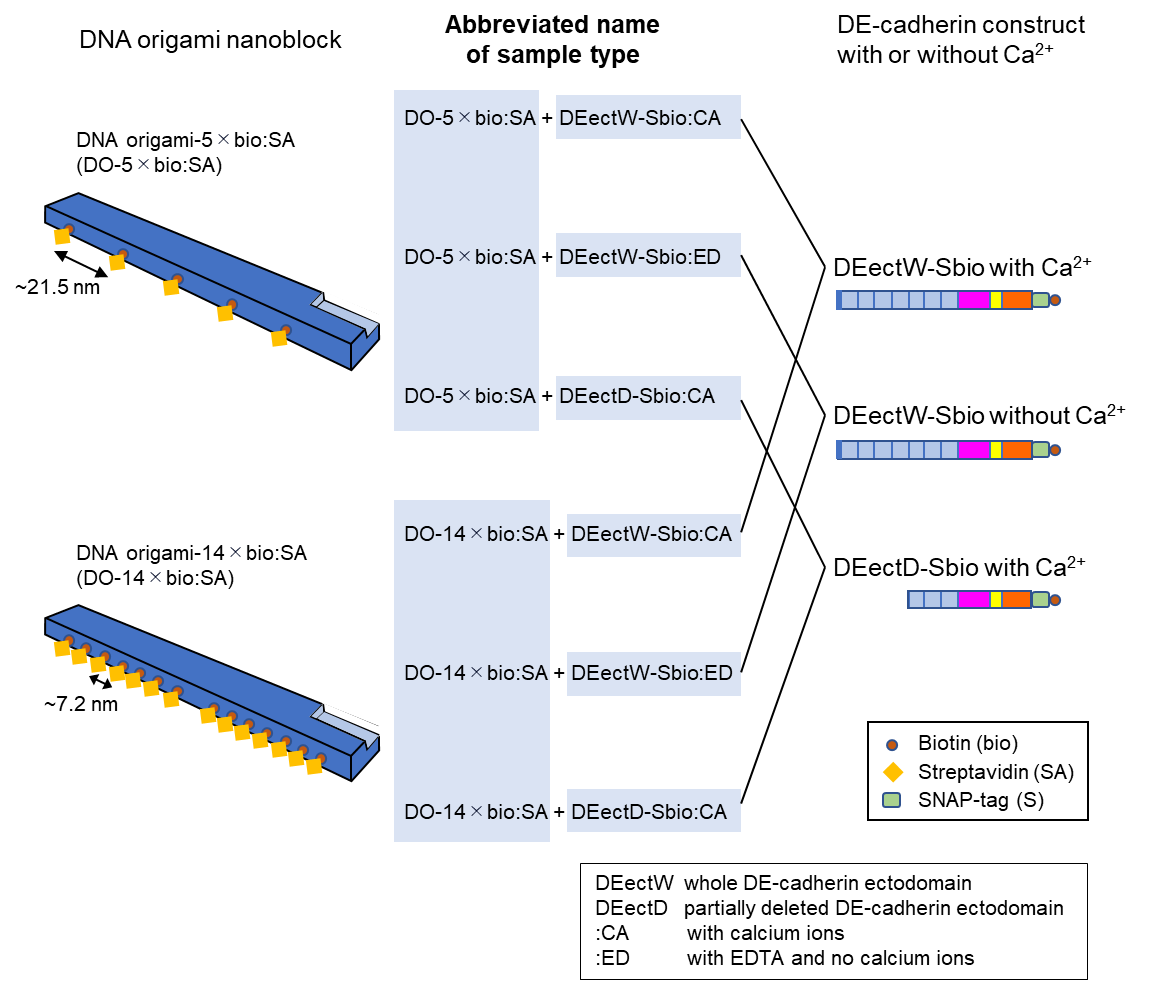

**Figure S4. Schematic representation of the naming system of sample types comprising the DNA origami and DE-cadherin construct.** Key graphic elements and abbreviations are explained in insets.

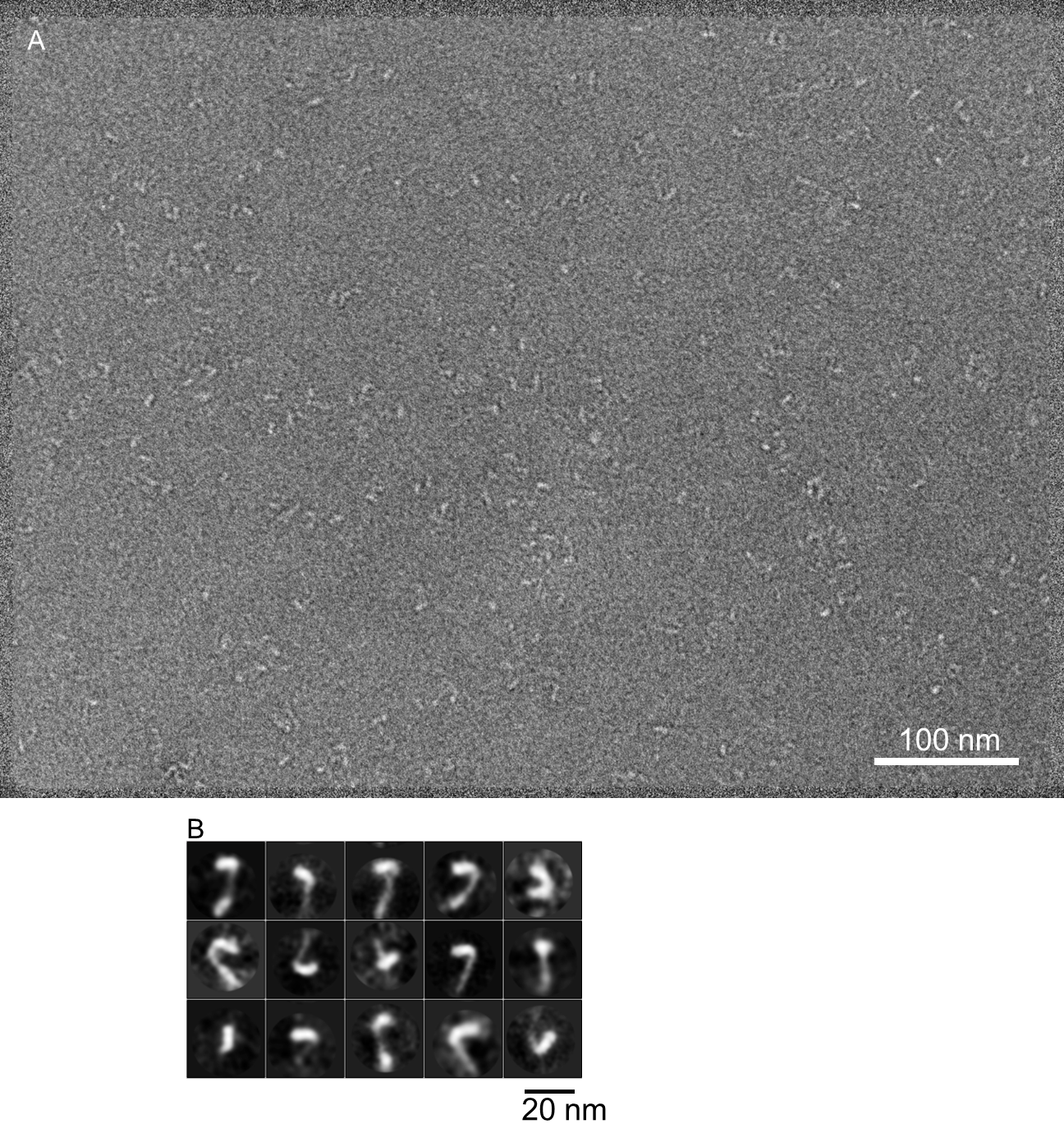

**Figure S5. Transmission electron microscopy (TEM) analysis of DEectW molecules.** (A) A TEM image of negative-stained, DEectW molecules. (B) 2-dimensional classification using 1,655 particles. Scale bars; 100 nm (A), 20 nm (B)

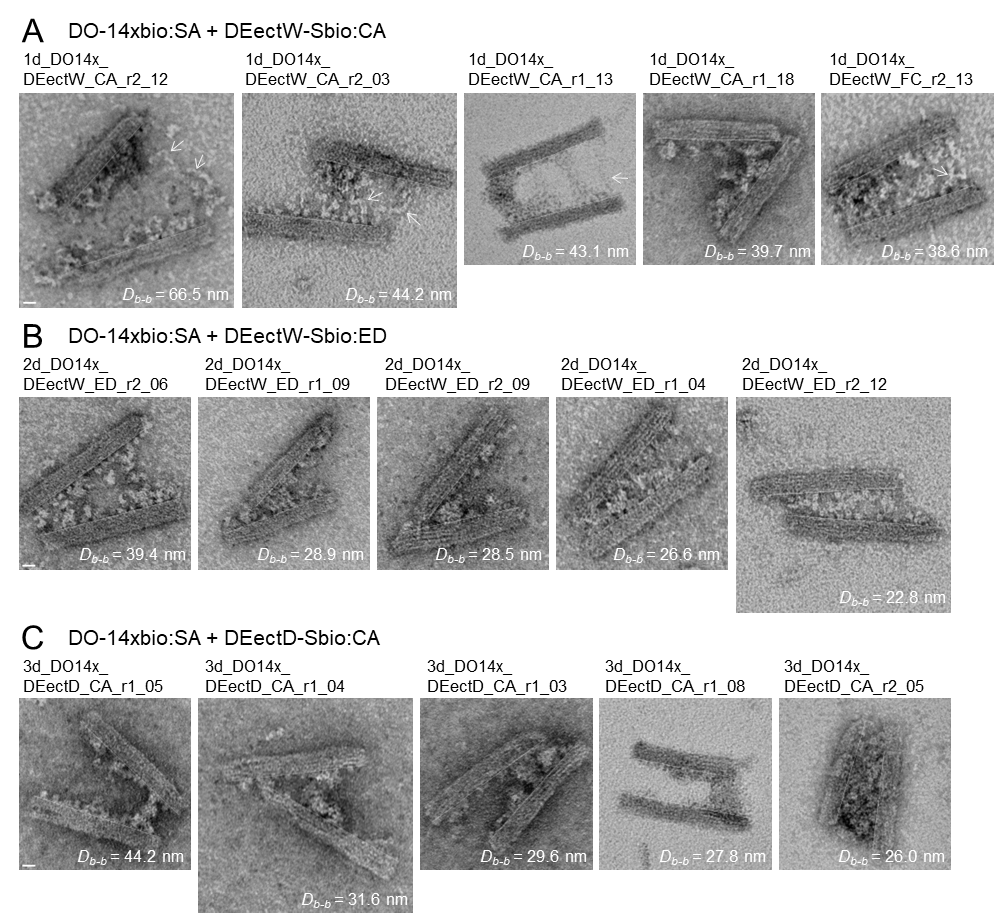

**Figure S6. Extreme examples of paired DO-14×bio:SA nanoblocks.** (A–C) Transmission electron microscopy images of paired DO-14×bio:SA nanoblocks having the five largest *D_b-b_* values, from each of the three sample types related to Fig. 5. (A) DO-14×bio:SA + DEectW-Sbio:CA. Arrows indicate string-like configurations of *Drosophila* E (DE)-cadherin ectodomain fragments contributing to bridging the apposed DNA origami nanoblocks. (B) DO-14×bio:SA + DEectW-Sbio:ED. (C) DO-14×bio:SA + DEectD-Sbio:CA.

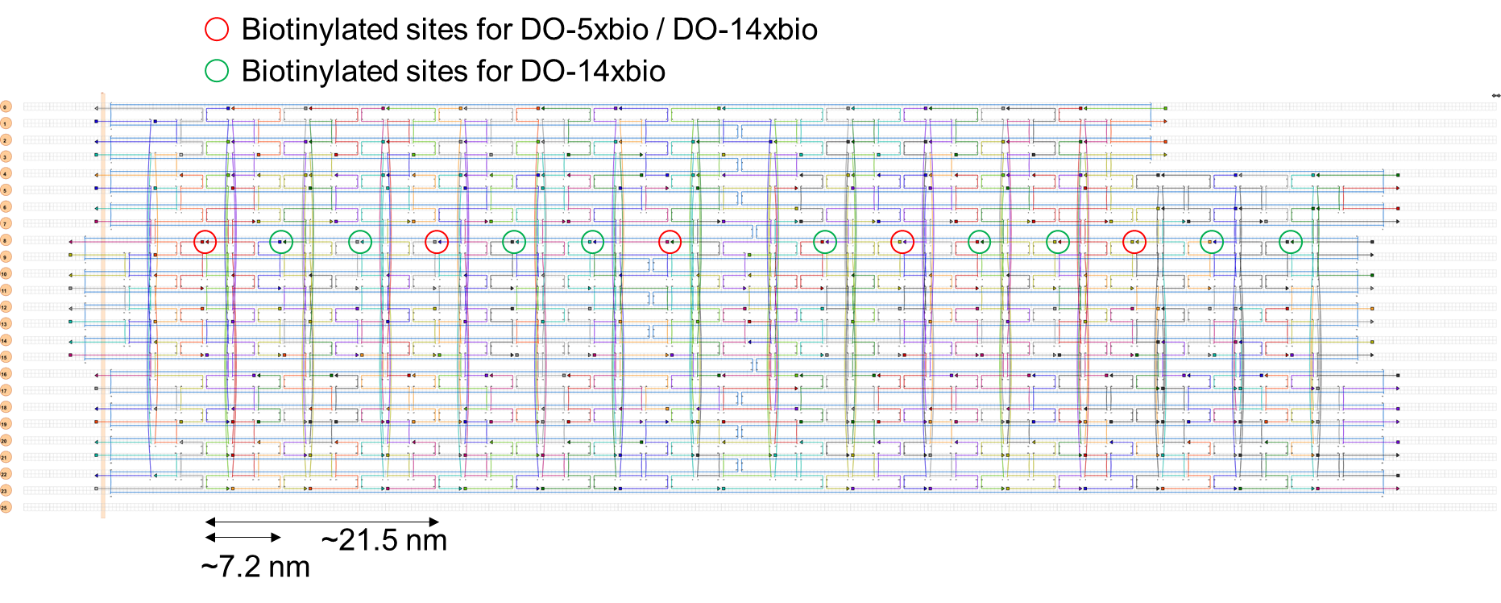

**Figure S7. DNA origami block design.** Annealing patterns of scaffold (p8064; light blue) and staple DNAs (239 species; arbitrary multiple colors) are shown. Positions of biotinylated sites for DO-5×bio and DO-14×bio are indicated. See Tables S2 and S3 for staple DNA details.

**Movie S1. Extending and shrinking motion of the membrane-distal portion of DEectW-Sbio on DO-5×bio.** The images were captured at a scanning speed of 0.3 s/frame. Image size: 60×60 nm^2^ (60×60 pixels).

**Movie S2. Binding and unbinding of neighboring DEectW-Sbio on DO-5×bio.** The images were captured at a scanning speed of 0.3 s/frame. Image size: 60×60 nm^2^ (60×60 pixels).

**Movie S3. Flexible motion of dimer-like structure formed between DEectW-Sbio on DO-5×bio and DO-5×bio-free DEectW-Sbio.** The images were captured at a scanning speed of 0.3 s/frame. Image size: 60×60 nm^2^ (60×60 pixels).
